# Human iPSC Derived Enamel Organoid Guided by Single-Cell Atlas of Human Tooth Development

**DOI:** 10.1101/2022.08.09.503399

**Authors:** A Alghadeer, S Hanson-Drury, D Ehnes, YT Zhao, AP Patni, D O’Day, CH Spurrell, AA Gogate, A Phal, H Zhang, A Devi, Y Wang, L Starita, D Doherty, I Glass, J Shendure, D Baker, MC Regier, J Mathieu, H Ruohola-Baker

## Abstract

Tooth enamel secreted by ameloblasts is the hardest material in the human body, acting as a shield protecting the teeth. However, the enamel is gradually damaged or partially lost in over 90% of adults and cannot be regenerated due to a lack of ameloblasts in erupted teeth. Here we use sci-RNA-seq to establish a spatiotemporal single-cell atlas for the developing human tooth and identify regulatory mechanisms controlling the differentiation process of human ameloblasts. We reveal key signaling pathways involved between the support cells and ameloblasts during fetal development and recapitulate those findings in a novel human ameloblast *in vitro* differentiation from iPSCs. We furthermore develop a mineralizing enamel organ-like 3D organoid system. These studies pave the way for future regenerative dentistry and therapies toward genetic diseases affecting enamel formation.

## Introduction

Tooth enamel is the hardest tissue in the human body. In addition to providing masticatory function, it protects the underlying dentin and dental pulp from mechanical, chemical, and microbiological damages that can lead to tooth loss. Unlike many other tissues, the adult human tooth does not regenerate enamel due to the absence of the enamel-secreting cell type, ameloblasts (Park et al., 2013), making enamel vulnerable to permanent damage or tooth loss. In addition to injury and damage, congenital genetic diseases such as Amelogenesis Imperfecta can also contribute to enamel loss. Ameloblasts are dental epithelial cells that secrete enamel protein matrix and deposit minerals to achieve hard and mature tooth enamel during human development (Jernvall and Thesleff, 2012). During tooth eruption in humans, ameloblasts undergo apoptosis (Park *et al*., 2013; Yajima-Himuro et al., 2014). Though almost all humans acquire some damage to the protective enamel shield as adulthood progresses, we currently do not have a way to regenerate ameloblasts (Fugolin and Pfeifer, 2017).

Although tooth development has been studied over several years (Yu and Klein, 2020), most of these excellent developmental and molecular studies have been conducted using murine models (Balic and Thesleff, 2015; Chiba et al., 2020; Krivanek et al., 2020; Sharir et al., 2019; Thesleff, 2014) which presents several challenges when applied to human development (Balic, 2019; Fresia et al., 2021; Hovorakova et al., 2018). For example, mouse incisors undergo continuous regeneration due to a population of epithelial stem cells in the labial cervical loop that allows for continued enamel formation throughout life (Harada et al., 2002). Since this regenerative process does not occur in adult human teeth, it is critical to understand tooth differentiation during early human developmental stages. In addition, the enamel organ, which ultimately gives rise to ameloblasts, is comprised of multiple populations of support cells, including the stellate reticulum and the inner and outer enamel epithelium (Nanci and TenCate, 2018). These support cells are thought to be essential for ameloblast function (Harada et al., 2006; Maas and Bei, 1997; Nakamura et al., 1991); however, it is not understood how they are mechanistically involved in ameloblast differentiation and functional maturation. Animal studies have suggested several pathways in driving and regulating this communication, such as the hedgehog (HH)(Koyama et al., 2001), NOTCH (Harada *et al*., 2006), and FGF (Takamori et al., 2008) pathways. However, the temporal regulation and the extent to which these pathways originate from support cells are not clearly understood since these cells are poorly studied in humans. Dissecting human tooth development at the single-cell level can capture the patterns of gene expression that characterize small populations of support cells that are involved in the differentiation.

In order to understand human tooth development and to facilitate the regeneration of human tooth structures in the future, we have utilized single-cell combinatorial indexing RNA sequencing (sci-RNA-seq)(Cao et al., 2019) technology to study human fetal tooth development at 9-22 gestational weeks (gw). Through computational analysis of the sci-RNA-seq data, we established for the first time a spatiotemporal single-cell atlas for developing human teeth that includes both the epithelial and mesenchymal cell types. Our computational studies established human-specific transcriptional profiles for subtypes of the developing tooth and revealed novel branches in the developmental trajectories of both mesenchymal and epithelial-derived tissues, as well as previously undescribed populations of epithelial support tissues. From our studies, we were able to identify, for the first time in developing human tissue, the subodontoblast, a proposed novel odontoblast progenitor. Further, we defined and induced the critical signaling pathways that drove changes in cell fate along the developmental trajectory of ameloblasts. This expedited the development of a 3D organoid that exhibits ameloblasts polarized towards odontoblast-like cells. This 3D organoid shows mineralization (calcium deposition) and expression of Ameloblastin, Amelogenin and Enamelin. Hence, we have coined the term Enamel Organoid to describe this new class of organoids.

These studies enhance our understanding of the regulatory mechanism controlling the differentiation process of dental tissues and lay the groundwork toward the development of disease models and regenerative approaches.

## Results

### A single-cell atlas of the developing human fetal odontogenic tissues

In humans, oral tissue development begins around 6gw and starts as a thickening in the oral epithelium (de Paula et al., 2017; Jussila and Thesleff, 2012; Nanci and TenCate, 2018), giving rise to all primary teeth and salivary gland tissue. Individual teeth develop independently as an extension of the main dental lamina and progress through a series of morphological stages (bud, cap, & bell) within bony crypts of the jaws (Radlanski et al., 2016). Additionally, each developing tooth is surrounded by thick fibrous tissue called the dental follicle (Wise et al., 1998). The dental follicle and the tissue it contains comprise the toothgerm (Kardos and Hubbard, 1981) (Figure 1A). The oral epithelium will also give rise to the salivary glands (Figure 1A). Like teeth, salivary glands derive from the invagination of a thickened sheet of oral epithelium into the underlying mesenchyme, known as the initial bud stage (Cha, 2017) (Figure 1A).

**Figure 1.**
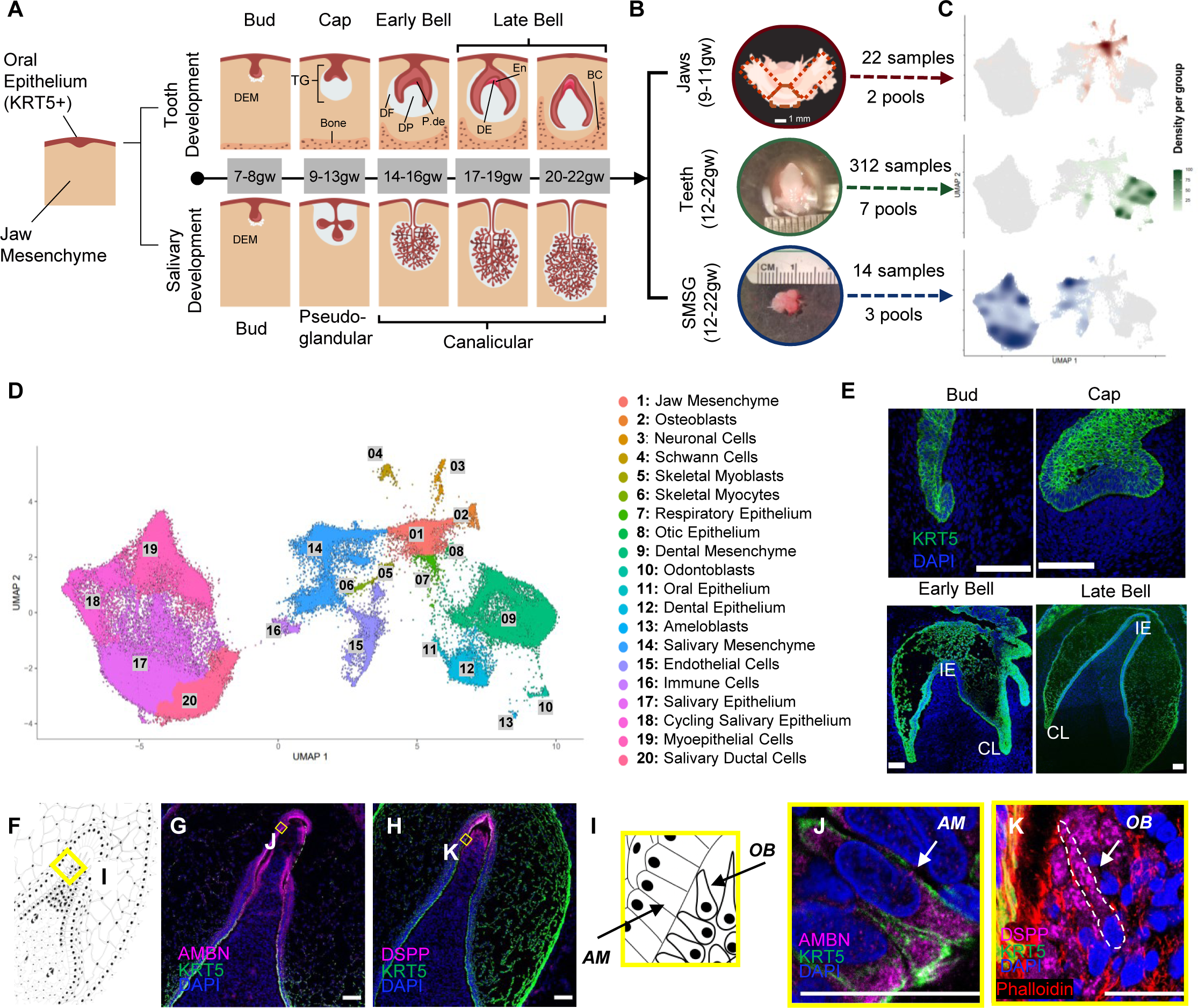
A single-cell atlas of the developing human fetal jaws, teeth, and salivary glands tissues *via* sci-RNA-seq. Human tooth and salivary gland exhibit stepwise developmental processes (A) The oral epithelium (colored in red) will give rise to the epithelial components of teeth and salivary glands, while the condensed dental ectomesenchyme (DEM, colored in grey) will give rise to the mesenchymal component of these tissues. TG: toothgerm, DF: dental follicle, DP: dental papilla, P-de: pre-dentin, De: dentin, En: enamel matrix. (B) Human fetal tooth germs and salivary glands were dissected in a stage-specific manner from human fetal jaw tissue. The young fetal jaws, 9-11 gestational weeks, were segmented into anterior segments (dotted box, which span from the canine-to-canine region) and posterior jaw segment pairs (dotted box) and sequenced independently. For older fetal jaws, 12-22 gestational weeks, individual toothgerms and submandibular salivary glands were dissected. A more detailed look at the dissection process can be found in Figure S1B. (C) Density plots of the clustered sci-RNA-seq data highlight the location of each tissue type in the same UMAP coordinate in D. The UMAP graph (D) yielded 20 annotated clusters from all sequenced data. (E) Immunofluorescence staining of developing toothgerms tissue sections with anti-Krt5 (green) that specifically marks the dental epithelial morphology throughout the developmental stages. Counterstained with the nuclear staining DAPI (blue). Abbreviations: incisal edge (IE), cervical loop (CL). To establish expression of known odontoblast and ameloblast markers in our tissue, immunofluorescence was performed on human fetal toothgerm at 20gw using dentin sialophosphoprotein (DSPP) and ameloblastin (AMBN), respectively (G,H) higher magnification in (J,K). Simplified illustration in (F, I). As expected, ameloblasts marked by AMBN (G, J) (pink), and odontoblasts by DSPP (H, K) (pink). Scale bars: 50μm.

To better understand early oral differentiation and to dissect how the epithelial and mesenchymal cell lineages acquire the odontogenic competence, we analyzed the developmental gene expression profiles of human fetal stages by single-cell sequencing. Toothgerm and salivary gland samples were collected from five fetal age groups (Figure 1A–1B and S1A–S1C). These age groups represented the following developmental stages for tooth differentiation: the cap stage (9-13gw), the early bell stage (14-16gw), and the late bell stage (17-22gw) (Figure 1A–1E) (Nanci and TenCate, 2018; Nelson, 2020). We also collected submandibular salivary glands (SMSG) from three matched timepoints (12-13gw, 14-16gw, 17-19gw) that cover the pseudo-glandular and canalicular stages for salivary gland development (Quirós-Terrón et al., 2019) (Figure 1A).

Single-cell sequencing data of the tissue samples were analyzed using Monocle3 (Cao *et al*., 2019; Trapnell et al., 2014) and visualized in uniform manifold approximation and projection (UMAP) space (Figure 1D). The distribution of the cells from each tissue origin was identified by using density plots based on tissue type (Figure 1C) or by individual samples (Figure S1D). Utilizing a graph-based clustering algorithm, we annotated 20 major clusters based on key marker genes (Figure 1D; Figure S1E; File S1) from PanglaoDB (Franzén et al., 2019). The major cell types in salivary gland samples include salivary mesenchyme, salivary epithelium, cycling salivary epithelium, myoepithelium, and ductal cells (Figures 1C–1D and S1E). In the jaw samples (9-11gw) (Figures 1C– 1D and S1E), we identified mesenchymal progenitors, osteoblasts, neuronal, Schwann cells, muscle, respiratory epithelium, otic epithelium, and oral epithelium (Figures 1C–1D and S1E). The major cell types in tooth samples include dental mesenchyme, epithelium, odontoblasts, and ameloblasts. The cell types observed in all samples include endothelial (Albelda et al., 1991; Jiang et al., 2016; Lampugnani et al., 1992) and immune (Böheim et al., 1987; Filion et al., 1990) cells. We have previously analyzed the salivary gland sci-RNA-seq data in more detail (Ehnes et al., 2022). The present manuscript focuses on the gene expression and signaling pathways governing tooth development.

To confirm the timing of the tooth morphological stages, we performed immunohistochemistry on tissue sections. As expected, all the enamel organ derived tissues were visualized by KRT5 (green) (Figure 1E). There are two critical lineages in tooth development: odontoblasts and ameloblasts. These two cell types secrete the mineralized protective layers that cover the soft dental pulp, which contains the nerves and the nutrient-transporting blood vessels. Odontoblasts are ectomesenchyme-derived cells secreting the inner coverage for the pulp, called dentin, while ameloblasts are ectoderm-derived and secrete the outermost layer, enamel. In order to establish the expression of known odontoblast and ameloblast markers in our tissue, immunohistochemistry was performed on human fetal toothgerm at 20gw using dentin sialophosphoprotein (DSPP) and ameloblastin (AMBN), respectively (Figures 1F–1K and S4K-S4Y). As expected, ameloblasts express AMBN in secretary vesicles (Figures 1I–1J); likewise, odontoblasts secrete DSPP (Figures 1I and 1K).

### Spatial localization of sci-RNA-seq defined clusters identifies subodontoblasts in humans for the first time and suggests they give rise to preodontoblast in an early developmental setting

To dissect the odontoblast lineage, we subset the developing jaw mesenchyme, dental ectomesenchyme, and odontoblast cells and embedded the data into a UMAP space (Figure 1D and Figure 2A). This analysis yielded six transcriptionally unique clusters: dental papilla (DP), preodontoblast (POB), odontoblast (OB), subodontoblast (SOB), dental ectomesenchyme (DEM), and dental follicle (DF) (Figure 2A). Cell types were identified by putative marker genes reported by monocle3 ‘top_marker’ function, which matches with recently identified odontoblast markers (Krivanek *et al*., 2020) (Figure 2B; Figure S2A–S2B; File S1 and File S2: meta-analytic estimators).

**Figure 2.**
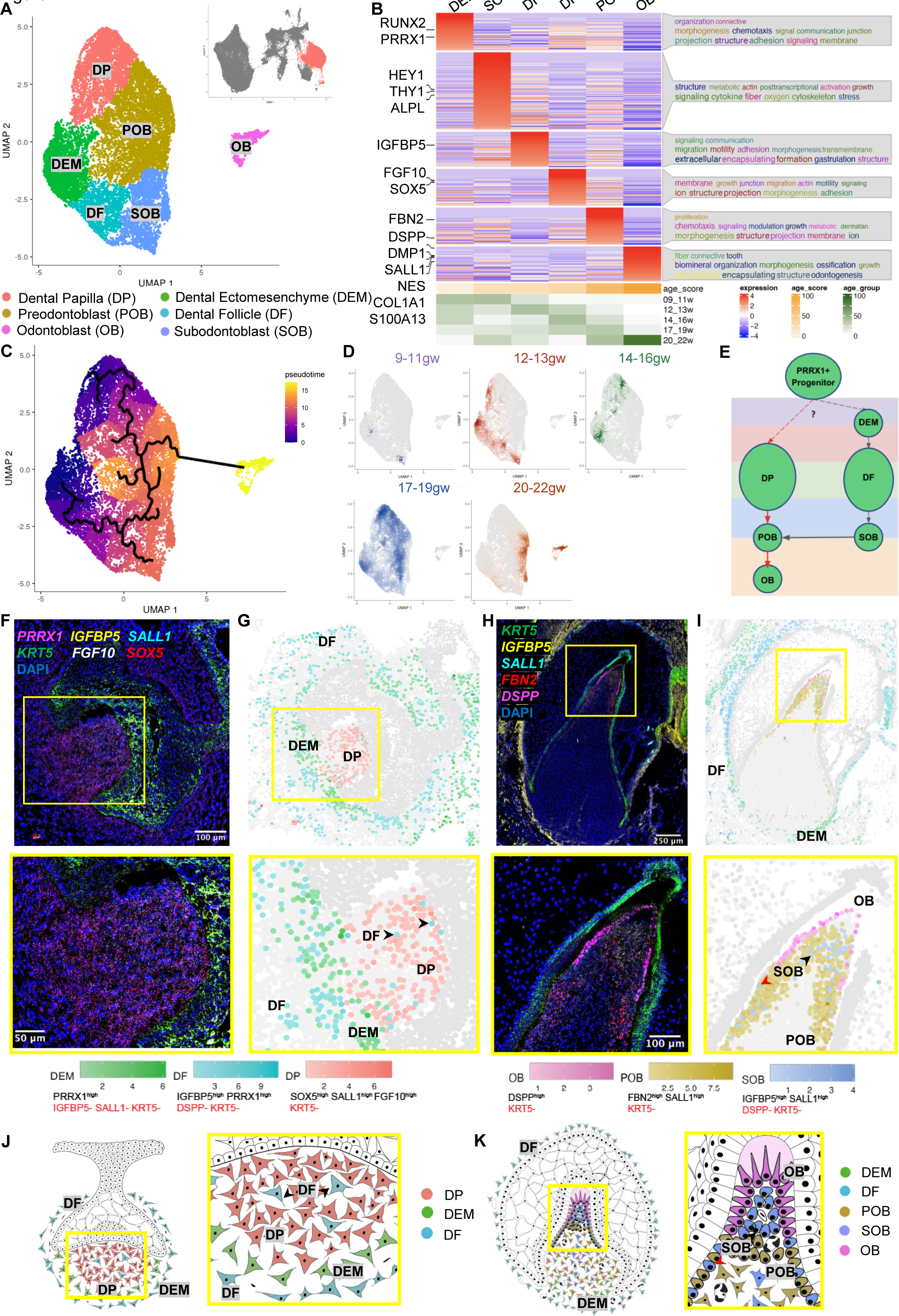
Dental Mesenchyme Developmental Trajectory. (A) UMAP graph of subclustered molar and incisor tooth germ type dental mesenchyme derived cells from the total dataset identified 6 transcriptionally unique clusters including dental papilla (DP), preodontoblast (POB), odontoblast (OB), subodontoblast (SOB), odontoblast (OB), dental ectomesenchyme (DEM), and dental follicle (DF). (B) A custom heatmap was generated to identify the marker genes specific to each cluster, the top associated GO-terms to characterize cluster function, and calculated age score per cluster. (C) Pseudotime trajectory analysis for dental mesenchyme derived cells suggest two progenitors DP and DEM (blue), that give rise to differentiated OB (yellow). (D) Real-time density plots indicate migration of cells from early progenitor populations DEM and DP at 9-16gw to differentiated DF, POB, SOB and OB at later development 17-22gw. (E) Simplified differentiation trajectory tree illustrating a common *PRRX1*+ progenitor gives rise to both DP and DEM. In the OB lineage (red), DP gives rise to POB, followed by OB, with a suggested SOB transitioning through POB-like state before giving rise to OB; and DF lineage (grey), of DEM giving rise to DF. (F) RNAScope HiPlex *in situ* hybridization image and inset including *PRRX1*, *SOX5*, *FGF10*, *SALL1*, *IGFBP5*, and *KRT5* probes and DAPI at 80d (G) RNAScope map for marker combinations corresponding to individual dental mesenchyme clusters for DEM (*PRRX1*+), DP (*SOX5+FGF10+SALL1+)* and DF (*IGFBP5+)* at 80d (black arrows indicate DF within the dental pulp) with individual cell types and stage matched replicates mapped in Figure S3. (H) RNAScope HiPlex *in situ* hybridization image and inset including *DSPP*, *IGFBP5*, *SALL1*, *FBN2*, and *KRT5* probes with DAPI nuclear stain at 117d (I) RNAScope map for marker combinations corresponding to individual dental mesenchyme clusters for OB (DSPP+), SOB (*IGFBP5*+*SALL1*+), POB (*FBN2*+*SALL1*+), and DF (*IGFBP5*+) at 117d (SOB beneath OB at incisal edge (black arrow) and intermingled with POB (red arrow)) with individual cell types and stage matched replicates mapped in Figure S3. (J) A diagram of the developing dental mesenchyme derived cell types of the human tooth germ. At cap stage (12-13gw) the dental pulp consists of DP cells with DEM with sparse DF within; DF surrounds the developing toothgerm. By bell stage (17-22gw), the dental pulp consists of OB at the incisal edge, SOB and POB with small contributions of the DEM and DP.

Furthermore, to evaluate the function of each cluster, we performed gene ontology analysis using ViSEAGO (Brionne et al., 2019), which uses data mining to establish semantic links between highly expressed genes in a given cluster. This analysis shows that DP and DEM are characterized by signaling, morphogenesis, and adhesion, supporting their role as precursor populations. In contrast, POB is characterized by their motility and migration, indicative of their alignment to the edge of the dental pulp. SOB indicates secretion, budding, projections, and branching, characteristics of a cell type sensing and influencing its environment, while OB shows GO-terms toward odontogenesis, tooth organization, and mineralization (Figure 2B).

To assess progenitor sources and cells’ progression towards differentiation, pseudotime trajectory analysis was performed. This analysis indicates the presence of two progenitor sources within the developing dental mesenchyme: the DP that gives rise to POB and, subsequently, OB; and the DF that gives rise to SOB, which transition through a POB phase before giving rise to OB (Figure 2C). Pseudotime analysis is supported by real-time density plots that show reduced progenitor type cell population density as the toothgerm develops, indicating fate commitment to OB lineage begins after 13gw in human fetal development and is largely complete by 20gw (Figure 2D). Broad expression of dental ectomesenchyme marker *PRRX1* is observed in both the DEM and DP (Figure S2C), supporting previous findings (Chai et al., 2000) that a shared cranial neural crest progenitor gives rise to both DP and DF. Thus, we propose a simplified trajectory of both the odontoblast and dental follicle lineages (red and grey arrowheads in Figure 2E), with a shared *PRRX1*+ progenitor giving rise to both DEM and DP.

To localize the computationally identified clusters in human fetal tissue, we performed RNAScope *in situ* hybridization on toothgerms at early (13gw, 80d) and late (19gw, 117d) tooth development (Figure 2F and 2H). After performing signal quantification per cell, we converted the RNAScope images into spatial datasets of single cells (Figure 2G and 2I; S2E–S2F). In agreement with the sci-RNA-seq data (Figure 2B; File S1; File S3– RNAScope_logic_tables.xlsx), dental mesenchyme-derived cell types display spatiotemporally specific expression patterns. At 13gw (Figures 2F–2G), the dental pulp consists of DP with DEM localized to the apical portion. The presence of sparse DF cells within the dental pulp (black arrowheads in Figures 2G and 2J) supports the pseudotime trajectory suggesting DF as progenitors for SOB, and this commitment occurs prior to 13gw. The developing toothgerm is surrounded by DF cells, a pattern that persists to late tooth development (19gw) (Figures 2H–2I). By 19gw, we observe that the dental pulp contains a mixed population of SOB and POB, with small contributions from DP, DEM, and OB at the incisal edge (Figures 2H–2I). We observe SOB directly beneath the OB (black arrowhead, Figure 2I) and, intriguingly, intermingled with POB at the pulpal periphery (red arrowhead, Figure 2I). This finding supports the pseudotime trajectory (Figure 2C), indicating SOB can give rise to OB not only following injury, as seen previously in mouse models (Harada et al., 2008; Ruch et al., 1995) but also during normal human tooth development. SOB represents a small portion of the pulpal cell population (Figure 2H–2I), suggesting that OBs are mainly derived from POB while SOB serves as a reserve with the capacity to differentiate to OB through a POB transitional state (Figure 2C; 2E and 2F–2K). This hypothesis is further strengthened by cell cycle analysis indicating SOB as a progenitor source of OB during normal tooth development, as this cell type has the highest proportion of cells in the G2M/S phase (Figure S2D). Lineage tracing studies are necessary to validate this exciting finding *in vivo* and further dissect SOB’s role in odontoblast development and repair.

### sci-RNA-seq and spatial localization reveal stage-specific support cell types and cervical loop stem cells for ameloblast differentiation

To further analyze the subtypes of the dental epithelium, we subset oral epithelium, dental epithelium, and ameloblast clusters (Figure 3A). The subset yielded 13 unique clusters that we identified by collating highly expressed cluster-specific genes (Figure 3B and S3A; File S1 and File S2: meta-analytic estimators). We were able to identify oral epithelium (OE), dental epithelium (DE), enamel knot (EK), enamel epithelium (outer enamel epithelium/inner enamel epithelium, OEE/IEE), cervical loop (CL), inner and outer stratum intermedium (SII, SIO), inner and outer stellate reticulum (SRI and SRO), pre-ameloblasts (PA) and two *AMBN* expressing ameloblast clusters (early ‘eAM’ and secretory ‘sAM’; Figure 3A–3D; File S1). The identity of these clusters aligned with their likely real-time appearance as represented by a real-time distribution of cells (Figure S3B). Moreover, GO analysis (Figure 3B) indicated cell type-specific roles in tooth development in agreement with our annotations. For example, the OE cluster revealed proper stratified epithelium, including keratinization, keratinocyte differentiation, and cornification (Adams, 1976), while the DE shows epithelial organization and differentiation, indicative of its function in reorganizing to form the tooth bud (Ahtiainen et al., 2016).

**Figure 3.**
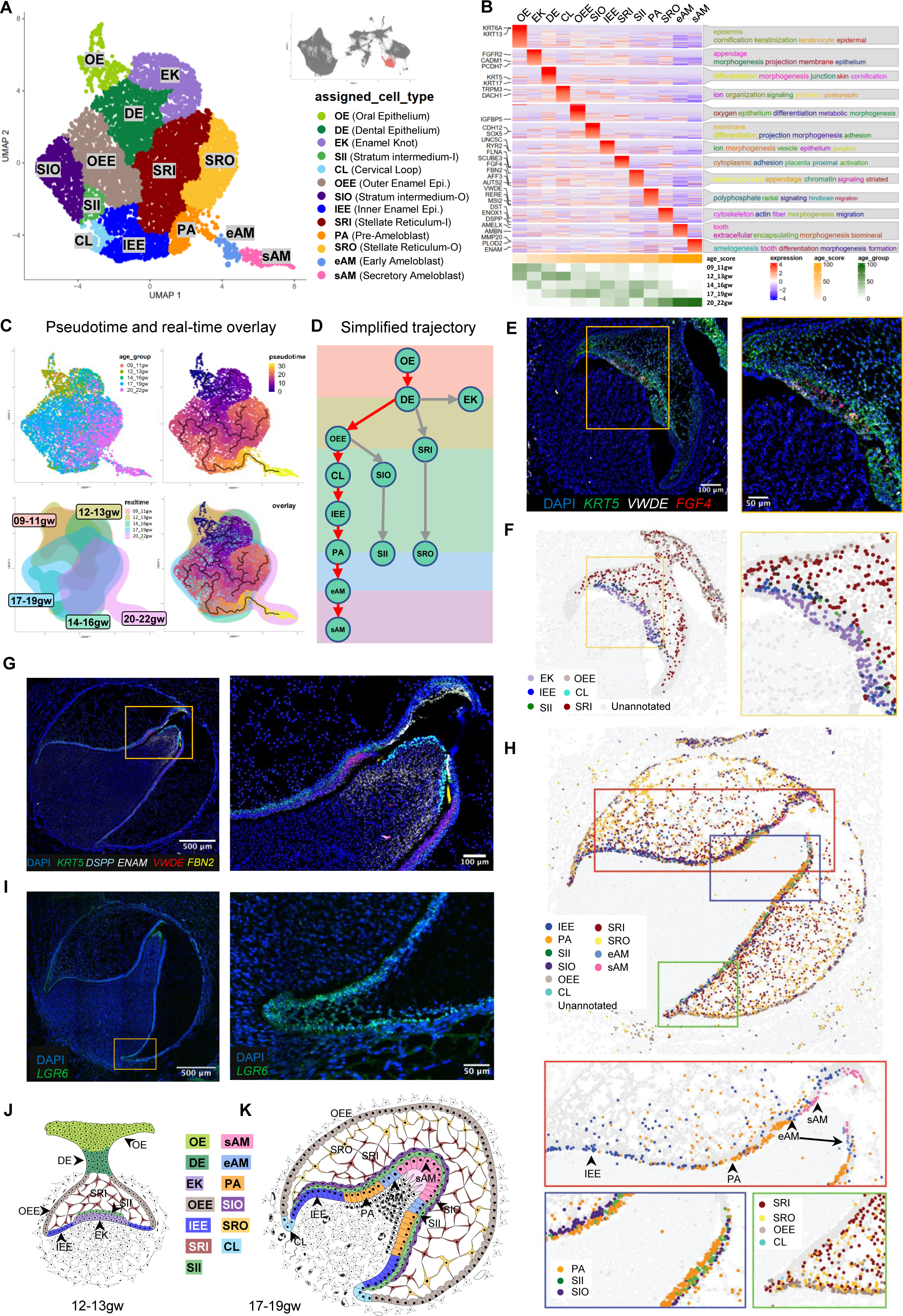
Ameloblast Developmental Trajectory. (A) UMAP graph of subclustered molar and incisor toothgerm type dental epithelium derived cells from the total dataset identified 13 transcriptionally unique clusters including the oral epithelium (OE), dental epithelium progenitors (DE), enamel knot (EK), outer enamel epithelium (OEE), inner enamel epithelium (IEE), cervical loop (CL), inner stratum intermedium (SII), outer stratum intermedium (SIO), inner stellate reticulum (SR), inner stellate reticulum (SRI), pre-ameloblasts (PA), early ameloblasts s(eAM) and ameloblasts (sAM). (B) A custom heatmap was generated to identify the marker genes specific to each cluster, the top associated GO-terms to characterize cluster function, and calculated Age Score per cluster. (C) Pseudotime trajectory analysis and realtime overlay for dental epithelium derived cells suggests that the DE directly gives rise to three branch lineages including the OEE, SR and EK. (D) Simplified differentiation trajectory tree illustrating the separate lineages originating from the DE, including the main AM lineage (red), of OEE, which gives rise to CL, IEE and PA, which then gives rise to eAM and sAM; and support cell trajectories (grey). (E) RNAScope HiPlex *in situ* hybridization image and inset for VWDE (high in IEE, SII, SRI), and FGF4 (high in EK) probes with DAPI nuclear stain at 80d (F) RNAScope map of individual dental epithelium-derived clusters - EK, OEE, IEE, CL, SII, and SRI – present at 80d shown as determined by relative expression of markers as specified in (File S3) with individual cell types and stage matched replicates mapped in Figure S6. (G) RNAScope HiPlex *in situ* hybridization image and inset for *DSPP* (high in eAM), *ENAM* (high in sAM), *VWDE* (high in SII, CL, PA), *FBN2* (high in IEE, SII, CL) probes with DAPI nuclear stain at 117d (H). RNAScope map of individual dental epithelium derived clusters – IEE, PA, SII, SIO, OEE, CL, SRI, SRO, eAM, and sAM – present at 117d shown as determined by relative expression of markers as specified in (File S3) with stage matched replicates mapped in Figure S3. (J) A diagram of the developing dental epithelium derived cell types of a toothgerm at 12-13 gestational weeks. The OE is lining the oral cavity while DE is the stalk connecting OE to the enamel organ. DE has given rise to the signaling center EK and SRI. OEE present at the periphery of the enamel organ have given rise to CL, SII and IEE. (K) A diagram of the developing dental epithelium derived cell types of a toothgerm at 17-19 gestational weeks. SII give rise to SIO layer, and together represent the superficial layer above IEE, PA, eAM and sAM, while SRI and SRO represent the bulk of the cells inside the enamel organ.

To identify the developmental trajectory of the dental epithelial lineages, we performed pseudotime analysis (Figure 3C) summarized by the simplified tree graphs (Figure 3D, ameloblast trajectory with red arrows). The trajectory analysis suggests that the OE directly gives rise to DE. The DE then gives rise to the EK and SR lineages and the OEE lineage, which gives rise to SI, IEE/PA, and eAM/sAM. In order to validate our bioinformatic findings, we performed RNAScope *in situ* hybridization at multiple timepoints. We used combinations of cluster-specific markers identified by transcriptional analysis to map cells from each cluster in the fetal tissues (Figure 3E, 3G, and 3I; Figure S3C; File S3– RNAScope_logic_tables.xlsx). Computational pseudo-spatial mapping of these cells revealed the following insights on EK, support cells, and CL function (Figure 3F and 3H).

The EK is a structure that has previously been identified at various times in mouse tooth development and is thought to organize local cell proliferation for epithelial budding or folding during cap and bell stage transitions (Thesleff et al., 2001; Vaahtokari et al., 1996; Yu et al., 2020). Primary EK has been shown to appear at the time of the first folding of the toothgerm to form the cusp, followed by secondary EK formation for subsequent cusp development. We identified a cluster of cells consistent with EK in human fetal development. Real-time distribution showed that cells occupying this cluster appeared at 9-11gw (early cap stage) and again at 14-16gw (early bell stage) (Figures 3B; S3B and S3D; 3E–3F), in line with the expected appearance of primary and secondary EK, respectively. EK are essential signaling centers in these early stages of tooth morphogenesis, playing a role in determining crown shape. Accordingly, GO terms identified in response to top gene expression associated with these clusters included morphogenesis and appendage development. These findings represent the first time this population has been identified at the transcriptional level and can lead to further understanding of the initiation of tooth morphogenesis and toothgerm type determination.

Multiple types of support tissues exist in the developing enamel organ. The SR are support cells with a star-shaped appearance in histological sections (Liu et al., 2016), which are thought to provide nutrients to and cushion the developing ameloblasts (Nanci and TenCate, 2018). Another support cell type, SI, is thought to support ameloblast differentiation (Liu *et al*., 2016)(Figures 3A–3B). We identified both types of support tissue in human fetal tissues. Furthermore, our single-cell analysis expanded upon what we understand about these populations. Transcriptomic analysis revealed two subgroups of SR, inner SR (SRI) closer to the inner surface of the toothgerm and outer SR (SRO) (Figures 3A–3D; 3H and S3A). Our analyses also identified, for the first time, two human SI sub-clusters that appear at 12gw and persist to later development (Figures 3A–3B; 3H and S3A). Inner SI (SII) represents the cell layer closer to ameloblasts lineage, and outer SI (SIO) represents the parallel support cell types adjacent to SII. The SI lineage at the early bell stage consists of two layers of cells, SII and SIO, that lie near ameloblast lineage (IEE, PA, and AM) (17-19gw) (Figures 3G–3H; 3K and S3E), creating a 3^rd^ previously unidentified stage-specific layer of cells (Figure 3H bottom left enlarged box). Furthermore, at the late bell stage, PA differentiates into eAM and matures to sAM (17-19gw) (Figure 3H top enlarged box, 3K). We propose these novel subgroups of support cells have precise signaling capacity to the specific, nearby epithelial cells in ameloblast lineage.

The enamel epithelium is the basal cell layer on the periphery of the tooth consisting of OEE, lining the outer side of the tooth, and IEE (Krivanek *et al*., 2020), lining the concave side of the folded tooth (Liu *et al*., 2016). As predicted, the transcriptional analysis revealed the presence of both of these populations, which was confirmed with RNAScope *in situ* hybridization (Figures 3A–B and 3H bottom right enlarged box; S3E). sci-RNA-Seq and RNAScope analysis revealed that in the cap stage, the core cells of the enamel organ are the DE that will give rise to the signaling center EK and the OEE (12-13gw) (Figures 3E–3F and 3J). sci-RNA-Seq also revealed a small population of LGR6+ CL cells expressing markers previously reported in epithelial stem cells of the regenerating adult mouse incisor (Chang et al., 2013). We could localize these cells in human fetal tissue to the expected location of the CL, where the OEE and IEE meet (Figures 3I and 3K). During the early bell stage, OEE are the basal cells on the periphery of the tooth organ that gives rise to SI, CL, & PA lineages (17-19gw) (Figures 3G; 3H and 3K). Importantly, the trajectory analysis predicts that the stem cells in CL can give rise to the ameloblast lineage. These data suggest that CL has a stage-specific role; while CL in humans is traditionally thought to be involved in later root development, our data suggest that during the early stage of fetal development, CL has a vital function in generating ameloblast lineage as the tooth crown expands. Further study is needed to determine these CL cells’ role in toothgerm type determination, including root number and morphology.

### Sci-RNA-seq reveals spatio-temporal expression patterns of critical signaling pathways in ameloblasts and facilitates the development of human iPSC-derived ameloblasts (iAM) *in vitro*

To understand the signaling pathways involved in ameloblast differentiation, we compiled a comprehensive multiplexed analysis pipeline based on ligand-receptor interactions and downstream transcriptional activity (Figure S5A). Briefly, a *talklr* (Wang, 2020) R package was used to identify specific ligand-receptor communications between the cell types at each developmental time point. DEsingle (Miao et al., 2018) and scMLnet (Cheng et al., 2021) programs were used to evaluate the downstream signaling activity by establishing multilayer networks between ligands and receptors and between transcription factors and their differentially expressed targets. Finally, activity scores were assigned to each pathway, which represent a percentage (0-100%) of the overall activity for all pathways included in the analysis.

Following our analysis pipeline, we evaluated the pathway activities between each stage in the ameloblast developmental trajectory and identified the most active pathways with specified ligands in each step (Figures 4A; S5A–S5B). To identify the main sources of the secreted ligands for each active pathway, we used *talklr* (Wang, 2020), with the ligand gene expression level analysis as a secondary validation (Figure 4B and S5C). These data revealed that during the transition from OE to DE, the BMP, ACTIVIN and noncanonical WNT (ncWNT) signals are secreted from the dental mesenchyme, while the canonical WNT ligands are secreted from within the OE.

**Figure 4.**
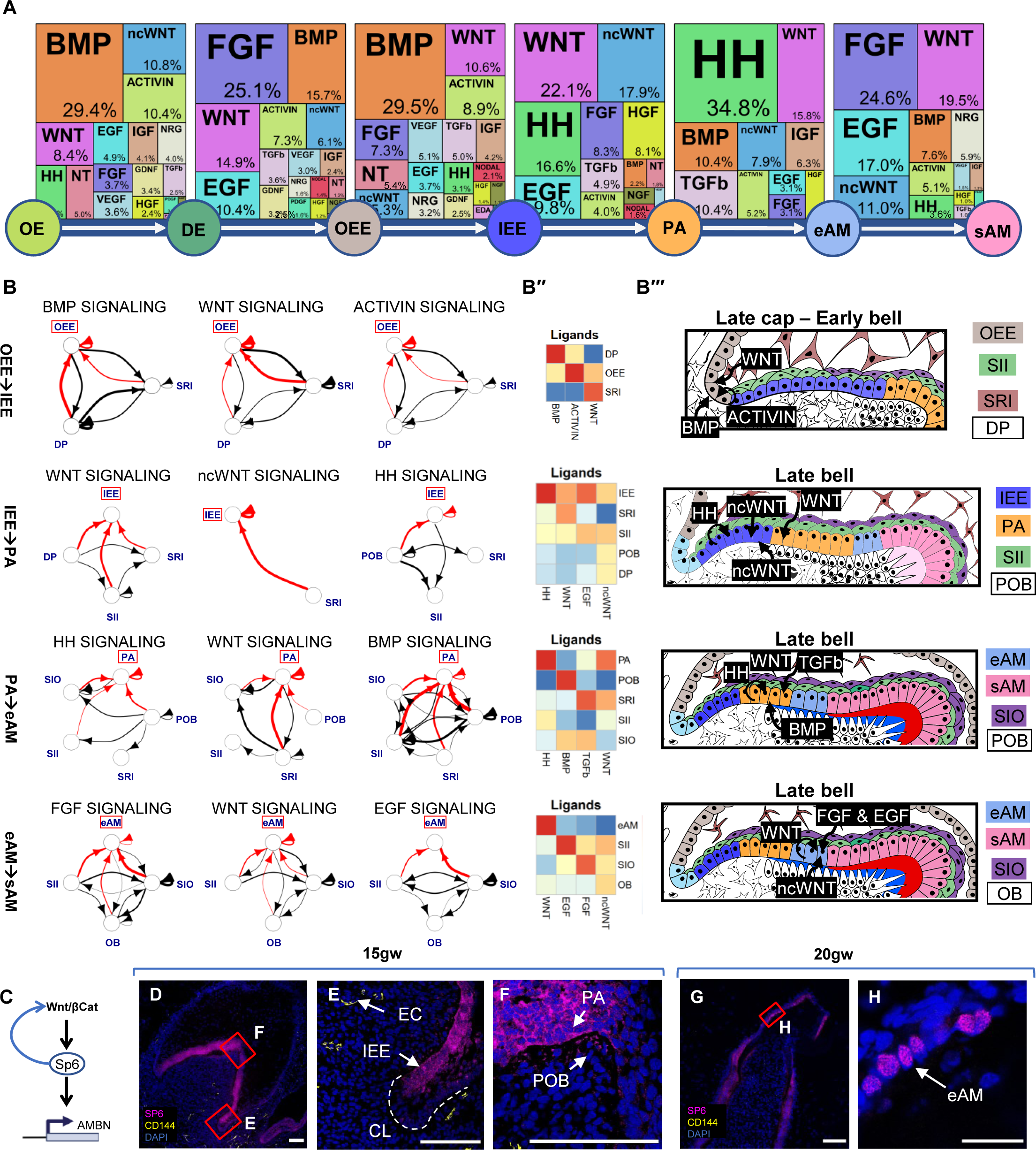
Signaling Pathway Analysis of The Ameloblast Trajectory. (A) The most active signaling pathways involved in ameloblast differentiation were identified to be BMP, WNT, HH and FGF, with detailed description of workflow found in Figure S5. (B) The sources of critical signaling ligands for the top three pathways involved for each developmental stage originate from both the dental epithelium and mesenchyme derived tissues, with the thickness of the line indicating the number of ligand:receptor interactions, arrowheads indicating the cell possessing the receptor, and interactions of interest (red) and between support cells (black). (B′′) Heatmaps for the top three or four pathways were generated by aggregating pathway ligand gene expression, which is then averaged per cluster. (B′′′) Diagrams illustrate the suggested ligand sources for each pathway at varying stages of tooth development. (C) Diagram for the proposed involvement of WNT pathway in activating the expression of SP6 which subsequently activate AMBN expression. Immunofluorescence staining of SP6 in 15gw tooth germ (D-F) confirm the start of expression of SP6 in cytosol of IEE where we predict the initiation of WNT activity in B’’’ at (OEE➔IEE), in the region of CL. Endothelial cells (EC) are present within the developing dental pulp at the same stage. At 20gw, SP6 is mainly localized to the nuclei coinciding with the onset of AMBN expression in differentiated AM at the tip of the toothgerm (G-H). Scale bars: 50μm.

Similarly, during DE to OEE, the signaling ligands are secreted from within DE and EK (Figure S5C). Meanwhile, the BMP and FGF ligands are mainly secreted from the surrounding dental mesenchyme. During the transition from OEE to IEE, the dental mesenchyme, which is now condensed as the dental papilla, mainly affects the ameloblast lineage by secreting BMP. Perhaps more interestingly, ligands for the prominent TGFβ pathway are mainly secreted from the support cells SRI, highlighting the importance of the spatio-temporal support cells in ameloblast differentiation. Other support types associated with stage-specific signaling behavior include SII ncWNT/HH/EGF, while SIO secretes FGF to support the last stages of ameloblast development and maturation. Additionally, mesoderm-derived POB and OB showed significant interaction with epithelial clusters; both secreted FGF and BMP ligands at PA during the PA to eAM transition or the transition to sAM. During ameloblast maturation, WNT ligands are mainly secreted from within eAM (Figure 4B).

Based on the signaling pathway prediction analysis, we found that BMP, ncWNT and ACTIVIN pathways are most active during OE to DE transition. However, ncWNT and ACTIVIN pathways down-regulate during DE to OEE transition when FGF and WNT pathways become more prominent. During the OEE to IEE stage transition, BMP is the most active, followed by WNT and ACTIVIN pathways. Meanwhile, in IEE to PA stage, mostly WNT (40% including canonical and non-canonical), HH (17%), and EGF (10%) become more active. In PA to eAM transition, the HH is the most active at (35%), followed by WNT, BMP and TGFβ. To analyze the maturation stage, we evaluated the pathway activities between eAM and sAM clusters and found that FGF, WNT and EGF signaling are involved in ameloblast maturation (Figures 4A and S5B). Our analysis of the stages of ameloblast development reveals a critical function for support cells, SI, SR and mesoderm signaling: BMP and ACTIVIN from mesenchyme are involved in the transition from OE to DE, ncWNT from DE, and support cells and BMP again from mesenchyme in the transition from DE to OEE, and from OEE to IEE. Similarly, IEE to PA differentiation utilizes specific accompanying SII to secrete ncWNT/EGF and WNT from SRI (Figures 4A–4B’; 4B’’’and S5B–S5C). In last stages of ameloblast differentiation, from PA to eAM, PA and SII secrete HH ligands, while SRI and SIO secrete TGFβ. At the final maturation stage, from eAM to sAM, SII secretes EGF & SIO secretes FGF. Interestingly, WNT activity in the transition of OEE to IEE (Figure 4B’’’) can be linked to the emergence of SP6 expression in IEE in the junction of the cervical loop (Figure 4C). WNT pathway has been suggested to work upstream of the expression of the transcription factor SP6 (Aurrekoetxea et al., 2016; Haro et al., 2014; Ibarretxe et al., 2012), which in turn was found to interact with AMBN/AMELX promoters (Rhodes et al., 2021) (Figures 4C–4F). Additionally, we found that SP6 is mostly localized in the cytoplasm of the early stages IEE/PA. However, in later stages, SP6 is localized to the nuclei coinciding with AMBN expression in eAM/sAM (Figure 4H). These data support the hypothesis that SP6 expression is induced by WNT pathway already in IEE transition stage, but becomes functional in eAM stage when it translocates to the nucleus and induces AMBN expression. Future loss-of-function analysis is required to test this hypothesis. Together these data suggest that WNT, TGFβ, HH, FGF, and BMP pathways are the top active pathways in ameloblast development compared to all 25 pathways included in the analysis.

We utilized the inferred signaling pathways from the sci-RNA-seq data (Figure 4A–4B) to develop a novel *in vitro* differentiation protocol that recapitulated the early stages of human ameloblast development from hiPSCs (iAM differentiation; Figure 5A). We first optimized a protocol to differentiate iPSC into OE (Ochiai et al., 2015; Suga et al., 2011; Tanaka et al., 2018). At day10 of differentiation, the OE markers were upregulated, while pluripotency markers were downregulated, and neuroepithelial and early mesodermal markers remained unchanged (Figure 5B: Figure S6B). To differentiate the OE cells into an early stage ameloblasts, we activated the main pathways we identified (Figure 4A) from the OE stage to the PA stage (BMP4, TGFβ1, WNT/CHIR99021, EGF, and HH/SAG pathways in the order of their activity during differentiation (Figure 5A; 5C). To transiently inhibit the BMP pathway, we used the small molecule LDN. This differentiation procedure resulted in marked epithelial morphological changes and a high expression of the ameloblast early marker *AMBN* at day 16 in the differentiation (Figure 5B; S6A–S6C), indicative of the early ameloblast differentiation stage.

**Figure 5.**
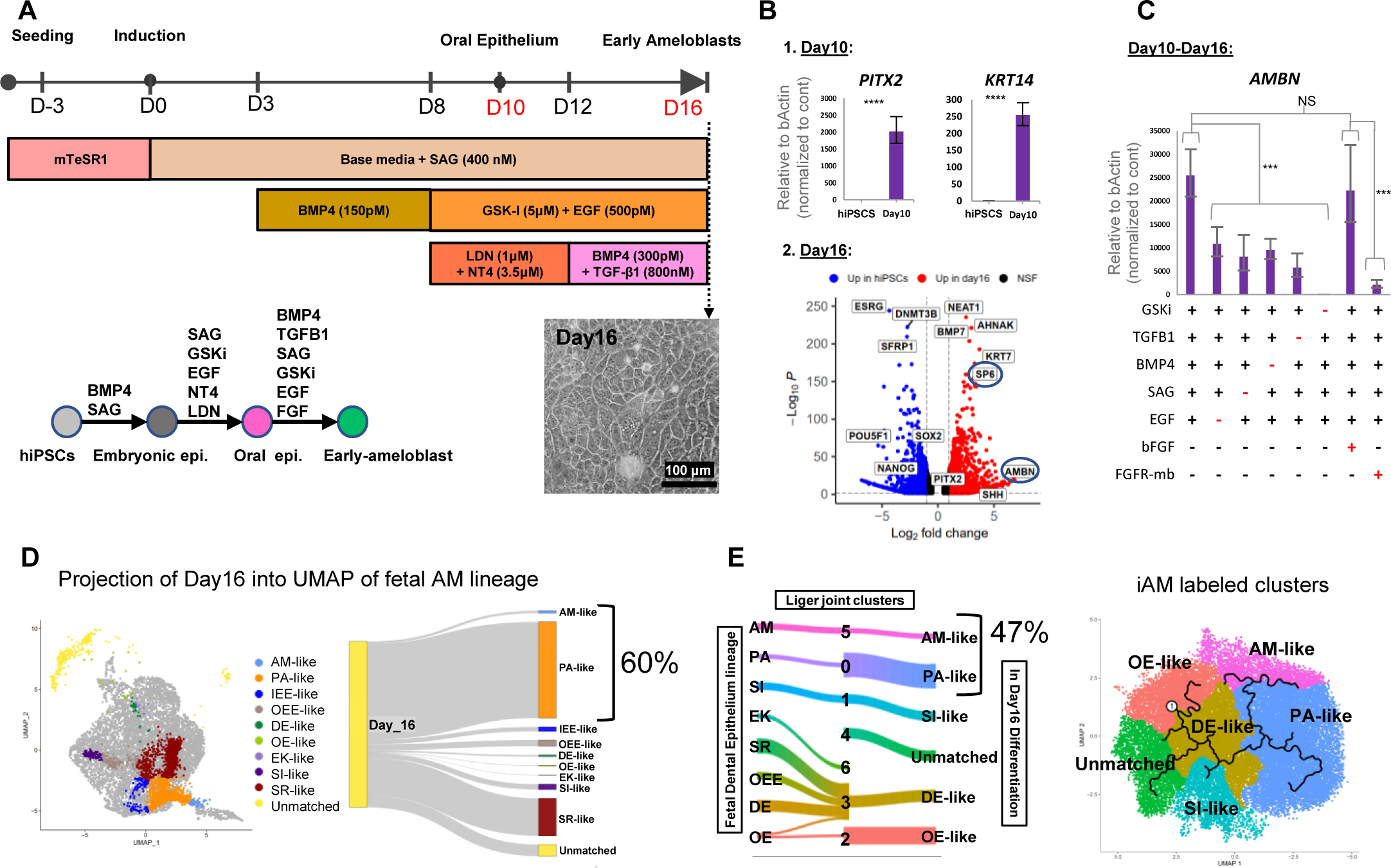
Human Induced Pluripotent Stem Cells (HiPSC) Derived Pre-Ameloblast Differentiation Protocol Guided by sci-RNA-seq. (A) Schematic of the 16-day differentiation protocol produced, which targets the identified signaling pathways utilizing growth factors and small molecules to transition through the ameloblast developmental trajectory. Cells at Day 10 of differentiation show upregulated expression of oral epithelium markers *PITX2* and *KRT14* as assessed by QRT-PCR (1B), while cells at Day 16 of differentiation show upregulation of ameloblast markers *SP6* and *AMBN* as assessed by bulk RNA-seq (2B) compared to undifferentiated hiPSC control. Each study was performed in triplicate, with error bars representing SEM. (C) The efficiency of each pathway activated in the differentiation from day10 to day16 were assessed by removing each agonist one at a time or adding FGFR2 mini binder to inhibit the FGFR2 pathway. *AMBN* expression was assessed in QRT-PCR, and each condition was performed in triplicate. Significance was determined by unpaired Student’s t-test; ***p□<□0.001; ****p□<□0.0001; Graph error bars are the means□±□SEM. (D) Projection of *in vivo* dental epithelium derived cell types (Figure 3A) with Day 16 identified clusters suggests 60% of Day 16 cells share gene expression pattern of PA and eAM (D) and LIGER joint clustering analysis suggests 47% of Day 16 cells share gene expression pattern of AM and PA (E), which allowed the labels to be transfer and annotate the cell types identity on iAM Day 16 differentiation UMAP graph.

To dissect which pathways were essential for the differentiation from day10 to day16, we used the process of elimination (Figure 5C). When removing EGF, SAG, BMP4, or TGFβ1 independently, we found that the expression of AMBN is significantly reduced to less than half compared to when all factors are present. However, removing GSKi completely abolished AMBN expression, suggesting that WNT signaling is a master regulator upstream to other pathways (Figure 5C). Our pathway prediction pipeline also suggested that the FGF pathway is heavily involved; however, adding bFGF had no significant effect on AMBN expression (Figure 5C). We hypothesized that the cells in culture secrete enough FGF ligands to saturate the receptors, and any exogenous ligands would have minimal effects. To test this hypothesis, we used a computationally designed protein, FGFR2 mini-binder (FGFR-mb), that specifically binds and inhibits the activity of the FGFR2 (Cao et al., 2022). Adding the FGFR2-binder almost totally abolished the expression of AMBN, which shows that the FGFR pathway is indeed required for ameloblast differentiation (Figure 5C). This marks the importance of highly specific AI-designed mini-proteins in analyzing the requirement of signaling pathways in differentiation. It is plausible that temporal preciseness and high penetrance, combined with specificity of designed mini-binders to their targets may partially out-compete in the future genetic perturbations of signaling pathway in iPSC derived differentiation paradigms.

To analyze the efficiency of the differentiation, we performed sci-RNA-seq on Day10 and Day16 of iPSC derived ameloblast differentiation (day10-OE and day16-Early-ameloblasts) and compared the gene expression data to the fetal tissue gene expression data. Our initial clustering and trajectory analysis indicated three major clusters at day10 and six clusters at day16 (Figure S6D–S6E). Sequencing revealed a significant overlap between human fetal and iPSC-derived ameloblasts in 2D culture. A survey of relevant markers to the dental epithelium (Figures S6F–S6G) showed the kinetics of their differential expression across the proposed trajectory (Figures S6D–S6E). Utilizing the markers for the oral/dental epithelial progenitors (Sun et al., 2016; Yu *et al*., 2020), enamel epithelium (Nakamura et al., 2017), and ameloblasts (Seidel et al., 2010), we were able to identify all the differentiated cell types (Figure S6E). For a better comparison between the *in vivo* and *in vitro* datasets, we used the projection method in Seurat 4.0 and the integration method in LIGER software packages (Hao et al., 2021; Welch et al., 2019) to overlay the datasets. We converted our dataset from Monocle3 format to Seurat format; then, we performed the projection over the UMAP of the fetal dental epithelium lineage. Lastly, we classified the projected cells using graph-based clustering. A small proportion of the cells in the day16 sample were OE-like, DE-like, SR-like, and SI-like. However, the majority (60%) were PA and AM-like, indicating that most of the differentiated cells are directed toward the ameloblast lineage (Figure 5D). We performed river plot analysis using LIGER to show the relationship between the annotated clusters from the fetal dental epithelial lineage and the *in vitro* day16 differentiation clusters that share the same space in LIGER joint clusters (Figure S6H), which allows label matching for the unannotated clusters in the differentiation (Figure 5E). Interestingly, the fetal OE cluster matched cluster 1 (d16_1; Figure S6E) in *in vitro* differentiation; the DE, SR, and OEE from *in vivo* samples mainly matched cluster 2 (d16_2) and SI matching cluster 4 (d16_4). The pre-ameloblast and ameloblast clusters matched clusters 5 and 6 (d16_5, d16_6), respectively, which represent 47% of total cells (Figure 5E). Finally, we analyzed the functionality of the iAM by analyzing the number of cells in day16 differentiated samples that produced AMBN, the product secreted by ameloblasts. Notably, 25% of the cells in 16 days of differentiation can produce and, in some cases, secrete AMBN protein (Figure S6C). This analysis suggests that our iAMs share similarities with fetal pre-ameloblasts and ameloblasts, demonstrating that the described 2D procedure can generate early differentiated ameloblasts.

### 3D Enamel organoids show mineralization and Ameloblastin, Amelogenin, and Enamelin secretion

To further characterize iAM and evaluate their capacity to mature *in vivo*, we injected the differentiated cells (day16, 2D) intramuscularly into adult SCID mice and allowed the injected cells to develop for 8 weeks (Figure 6A). The injected region was identified by human nuclear antigen staining (Figure 6B). The maturation stage of the iAM cells was analyzed in the subsequent serial sections by definitive ameloblast markers: AMELX, AMBN, DSPP, KRT14, and by its calcification capacity. Importantly, the identified iAM cells were significantly more mature (Figure 6B; 6E and S7A–S7G), showing that the iPSC derived iAM cells have a capacity to develop to a more mature AM stage. The highly elongated morphology of these cells (Figure 6D) suggests that they have developed into so-called secretory stage AM (sAM) that characteristically consists of tall columnar cells that express amelogenin (AMELX) and ameloblastin (AMBN) and produce mineralization. Accordingly, we further identified iAM capacity to produce calcified material *via* Alizarin red and Von Kossa staining (Figure 6E and S7G).

**Figure 6.**
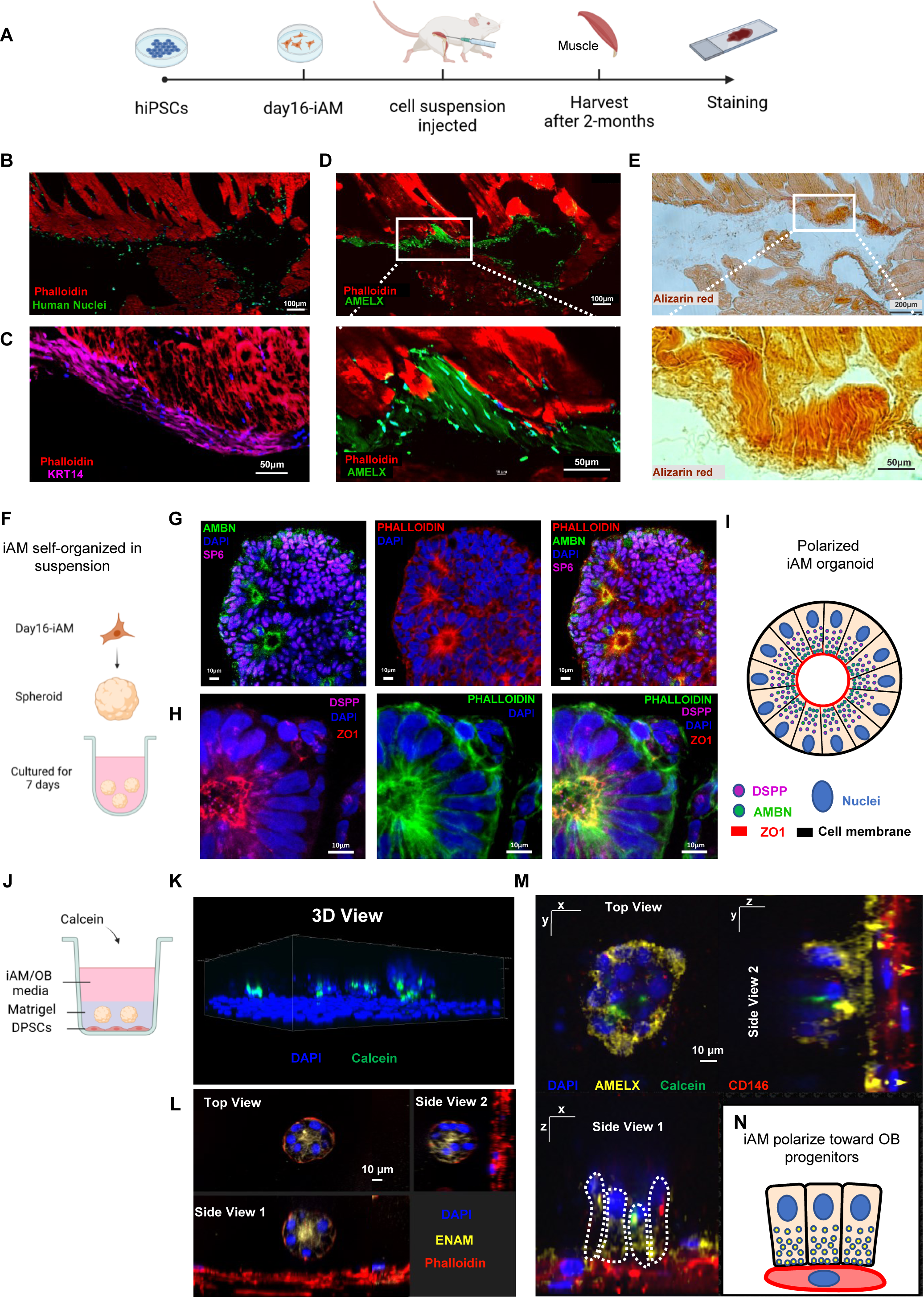
Characterization of iAM and formation of iAM/OB organoids. (A) Schematic of the mouse *in vivo* experiments describing the steps for injecting day16-iAM subcutaneously into the left legs muscle of the adult SCID mice. The adult SCID mice at 2-month-old were dissected at the site of injection to perform further analysis to locate the cells such as immunofluorescence staining for human nuclear antigen in (B), KRT14(C), AMELX (D) and Alizarin red staining in (E) showing mineralization. (F) Schematic of iAM organoids formation while cultured in suspension in ultra-low attachment plate. The formed iAM organoids express SP6 in the nuclei and secrete AMBN (G), and DSPP toward the apical side indicated by ZO1 (H). (I) A diagram simplifying the iAM organoid polarized structure toward a central lumen marked by ZO1, and the secretory vesicles of DSPP and AMBN. (J) Schematic for the coculture experiment between DPSCs as monolayer and iAM embedded in Matrigel above it. Calcein, which is a fluorescent dye that binds to calcium, was added to the media containing iAM base media and odontogenic media in 1:1 ratio. (K) 3D reconstructed image from z-stacked confocal images captured from the coculture experiment plate. The cocultured organoid show association with Calcein, as well as expression of ENAM at the center after 7 days (L), and after 14days the organoids close to CD146-expressing DPSC/OB, started to revert polarity towards DPSCs/OB while expressing AMELX (M) as simplified in the diagram in (N).

Since close contact between AM and OB is critical for tooth development, we proceeded towards developing an organoid model of the two cell types. We first developed an organoid model of polarized AM. To generate cells expressing AMBN in a culture with apical-basal polarization, similar to ameloblasts *in vivo*, we grew the cells in suspension to form spheroids (Figure 6F). We then performed immunofluorescence staining for SP6, AMBN (Figure 6G), and ZO1, DSPP (Figure 6H). We observed that the transcription factor SP6 is expressed in all the differentiated cells and is exclusively localized to the nucleus. The induced ameloblasts show apical-basal polarity and secrete AMBN to the apical surface. As seen in *in vivo* AM, the nucleus is located towards the basal side of the cell (Figure 6H–6I). Early ameloblasts are known to transiently express DSPP during development (Figure 1H; Figure 3G and S4X–S4Y); we noticed that DSPP expression is also localized to the apical side of the induced cells. Moreover, the tight junction protein ZO1 marks the apical side of these iAM cells (Figure 6H –6I). The iAM in the organoids appear as tall columnar cells polarized toward a central lumen (Figure 6I).

We next co-cultured the induced ameloblast organoids with primary human dental pulp stem cells (DPSCs) to assess the interaction level between the two cell types and the effects on ameloblast maturation. We found that simple coculture in suspension can induce AMELX in iAM organoids and DSPP in the odontoblast organoids, as observed in the developing human tooth (Figure S4A–S4J); as well as induction of calcified matrix (Figure S7H– S7K). After confirming that iAM can mature in the presence of OB/DPSCs, we designed the following experiment to coculture the cells in a layered approach. We plated the DPSCs in the bottom of a flat bottom plate and then embedded iAM organoids in a Matrigel layer above the DPSCs (Figure 6J). The co-culture media contained iAM and odontogenic media at 1:1 ratio, with calcein in addition to detect calcification. Through 3D reconstructed confocal images, we observed that iAM are associated with calcein, demonstrating the capacity of iAM to produce mineralization/calcification (Figure 6K). Furthermore, the co-cultured iAM expressed ENAM and AMELX (Figure 6L) and reverted their polarity towards the differentiating OB (Figure 6L–6N). This 3D organoid, therefore, mimics the normal cell-to-cell interface observed in developing tooth where the enamel proteins are secreted towards the OB and sets the stage towards developing human tooth organoids in a dish.

## Discussion

Functional ameloblasts and odontoblasts are two critical cell types secreting the protective tooth coverings, enamel, and dentin, that are required to generate the functional structure of teeth. Ameloblasts do not exist in adult oral structures, making enamel regeneration impossible; however, dentin-secreting odontoblasts are critical for the regeneration of adult teeth. While previous morphological studies have suggested that two cell types can give rise to odontoblasts, the developmental lineages and molecular characterization of this process were not understood. Here we generate and utilize single-cell sequencing to identify the cell types in the developing human tooth and their molecular interactions across several developmental stages. We identified the major cell types in human oral development that derive from the jaw tissue and give rise to teeth and salivary glands. Significantly, we show the presence of subodontoblast cells for the first time in human tissue. Additionally, we further characterized mesenchymal and epithelial odontogenic progenitors and revealed potential developmental trajectories that lead to odontoblasts and ameloblasts, respectively. Importantly, we identified novel human support cell types that significantly and precisely promote the differentiation of ameloblasts. Analyzing the signaling interaction in the ameloblast trajectory allowed us to predict the signaling molecules needed to recapitulate ameloblast development *in vitro*. Utilizing these findings, we developed a novel differentiation protocol to drive the differentiation of iPSCs toward early ameloblasts (iAM). We successfully verified their identity by comparing the expression profile of the *in vitro* generated ameloblast lineage to the *in vivo* fetal counterpart. Finally, we used this information to develop an enamel organoid that expresses mature ameloblast markers and secretes mineralized calcium.

The sci-RNA-seq data revealed novel transcriptionally defined subgroups of cells in both the epithelial and mesenchymal lineages. Our analyses identified 13 subclusters of cell types in the dental epithelial lineage, of which SRI, SRO, SIO, and SII are novel types of support cells in human tooth development. The newly identified support cell type, SRI, produces a TGFβ ligand at an early stage of tooth development to aid in the differentiation of IEE to PA. While SII secrete EGF and SIO secret FGF ligands at later stages to aid in the maturation of AM. Our data have amended the detail with which we understand how support cells contribute to the patterning and development of ameloblasts. Our analysis also revealed a previously undescribed role for SOB in the mesenchymal lineage. We show that while human OB are derived from POB, surprisingly, the developing human fetus has two potential sources that generate POB: DP or DF. Both precursor stages are strictly found in early fetal tissue; after 20gw, these precursors are largely absent in the dental pulp. A portion of DF cells differentiates to SOBs that have a more permanent, but previously unidentified function in tooth development. We now characterize this novel SOB cell type on a molecular level and show that they have characteristics of cells that sense and influence their environment, supporting the idea that these cells can sense the need to regenerate the lost OB population (Harada *et al*., 2008). Furthermore, a well-known disease gene in tooth development (Duverger and Morasso, 2018; Duverger et al., 2012), *DLX3*, is a key marker for SOB, calling for further analysis of *DLX3* function in this critical preserved group of cells in a disease-in-a-dish approach.

For the first time in human tooth development, our studies have revealed, in extreme detail, the signaling pathways that govern each transition between cell identity. Previous studies of hypodontia and tooth agenesis have shown that disruption of WNT, BMP, and FGF signals results in defective tooth development. However, the detail with which our study has revealed the role of these pathways at various points in development may more mechanistically explain how defects in these pathways lead to tooth loss or tooth agenesis. For example, studies have shown that mutations in BMP4 correlated to tooth agenesis (Yu et al., 2019a). Our analysis showed that BMP4 signaling is critical during the early stages of both the OE to DE and DE to OEE transition, suggesting that loss of BMP4 may lead to agenesis by disrupting these transitions. In other studies, disruption of FGF signaling leads to enamel irregularities (Marangoni et al., 2019). Using AI-based protein design, we now revealed that FGF signaling is essential at the point of ameloblast maturation, suggesting that these irregularities are a result of failure of ameloblasts to mature. While some studies have focused on the role of a single signaling pathway, many others have highlighted the importance of crosstalk between pathways in tooth development and maintenance (Liu et al., 2020; Malik et al., 2018; Yu et al., 2019b). Our predictive pathway analysis highlights not only the primary pathway responsible for each stage but also ranks the other pathways involved. Overall, our study will facilitate the investigation into both previously identified and yet undescribed crosstalk in driving forward development. The detailed analysis we have provided in this study will facilitate more detailed and informed studies on degenerative dental diseases and can lead to developing more effective ways to mitigate or reverse tooth loss. Furthermore, our work with AI-designed, de novo receptor mini-binders that specifically bind and inhibit target receptor signaling (Cao et al., 2022) now reveals a novel, highly simplified method to identify the exact stage of a specific signaling pathway required in the differentiation process. The method described in this study using the de novo FGFR-mb to unravel the FGFR pathway requirement in Ameloblast maturation will be generally applicable and specific to any signaling pathway analyzed in the differentiation of normal and disease organoids.

Ameloblasts secrete the most mineralized and highly vulnerable layer in the human tooth. However, this cell type, and hence enamel regeneration, is absent in adult humans, presenting an impasse for progress in human regenerative dentistry. Our studies have revealed multiple new potential avenues through which further study could overcome this. First, our studies present the first single-cell analysis and *in vivo* localization of the cervical loop in human fetal teeth. The cervical loop is part of the enamel organ in the developing tooth located where the OEE and the IEE join. It has been extensively studied in the mouse, most often in the mouse incisor. However, unlike the human tooth, the mouse incisor grows continuously, with the cervical loop serving as a reservoir of stem cells that contribute to that consistent growth. Therefore, it is necessary to understand the function and contribution of the cervical loop in human tissue. Classically, the cervical loop is known to give rise to Hertwig’s Epithelial Root Sheath, which initiates root formation. Intriguingly, our analysis revealed a role for the CL in giving rise to human ameloblasts in early tooth development, as the crown expands before the root begins to form. Our findings provide a basis for future studies to develop cervical loop-like cells with ameloblast lineage potential.

Finally, the present work characterized the molecular basis for human ameloblast differentiation. We have used this knowledge to develop an assay for differentiating human iPSC-derived ameloblasts in a dish (iAM). Comparing fetal data to the iAM differentiation suggests that iAM shares high similarity with fetal pre-ameloblasts and early ameloblasts. Further, iAM can reach the secretory stage since Ameloblastin protein production and secretion is observed in these cells. In addition, the iAM cells showed a significant increase in maturation, including calcifications, when tested *in vivo*. Upon co-culturing iAM and OB lineage we observed the iAM reverting their polarity and apical secretion of enamel proteins toward the OB lineage cells. Hence, we argue to have developed a chemically defined serum-free differentiation protocol to generate human dental epithelium, and their subsequent differentiation into enamel organ-like 3D organoids. This developed organoid has potential for future dental therapies.

The first human single-cell tooth development atlas described here paves the way toward successful human regenerative dentistry in the future. The molecular analysis and *in vitro* ameloblast differentiation protocol allow future dissection of diseases such as Amelogenesis Imperfecta that will guide the field toward therapeutic approaches.

## Materials and Methods

### Tissue collection and dissection

This study is approved by the Institutional Review Boards (IRB) at University of Washington for the use of human fetal tissues: BDRL (CR000000131) and Ruohola-Baker Laboratory (STUDY00005235). Fetal craniofacial tissues were collected from Birth Defect Research Laboratory (BDRL), University of Washington, and transferred to Ruohola-Baker laboratory submerged in Hank’s Balanced Salt Solution (HBSS) media (Gibco, #14025092) on ice. Toothgerms and salivary glands were dissected in cold RNase free Phosphate-Buffered Saline (PBS) (Invitrogen, #AM9624) within six hours from the initial dissection at BDRL. To extract the toothgerms, a vertical cut was made at the midline of the upper/lower jaw for orientation, then a horizontal cut was made from the right side of the midline along the top of the alveolar ridge to expose one toothgerm at a time. The first two toothgerms from the midline were the incisors, the next toothgerm was the canine, and the last two toothgerms were the molars. The same procedure was followed to extract toothgerms on the left side of the jaw. The submandibular salivary glands were harvested from the distal end of the lower jaw. The toothgerms from 9 to 11 weeks old were too small for dissection and not useable for sequencing; therefore, these jaws were cut into two posterior sections and one anterior section to separate molars from the incisors and canines at these timepoints. The extracted tissues were transferred into an Eppendorf tube and snap frozen using liquid nitrogen. The frozen samples were stored at –80°C until nuclei extraction.

### Nuclei extraction

Frozen tissues were carefully transferred to a stack of chilled aluminum foil kept on dry ice to prevent thawing. The folded foil encapsulating the tissues were placed on a block of dry ice and the foil was pounded with a pestle to pulverize the tissues into powder. 1mL of lysis buffer that contains nuclei buffer (10 mM Tris-HCl, 10 mM NaCl, 3 mM MgCl_2,_ pH 7.4), 0.1% IGEPAL CA-630, 1% SUPERase In RNase inhibitor (20 U/μL, Thermo), and 1% BSA (20 mg/mL, NEB) were added onto the tissue powder and transferred to a 1.5mL tubes. Samples were incubated in the lysis buffer for 1 hour on ice. The samples were pipetted up and down with pre-cut 1000uL pipette tip to disassociate the tissue further. The dissociated tissues were passed through 70 um cell strainers (Corning) into a 50mL conical tube. The strainers were rinsed with lysis buffer to minimize nuclei loss. The samples were centrifuged to pellet the nuclei at 500g for 5 minutes at 4°C and the supernatant was discarded. The samples resuspended again in 1ml lysis buffer, transferred into new 15mL tubes, pelleted again and the supernatant was discarded. The pellets were resuspended in 50ul of nuclei buffer, and 5 mL of 4% Paraformaldehyde (PFA) (EMS) diluted in RNase free PBS, was added to fix the nuclei for 15 minutes on ice. The tubes were flicked gently every 5 minutes to reduce clumping of nuclei. The fixed nuclei were pelleted at 500g for 3 minutes at 4°C and the PFA waste was discarded. The pelleted nuclei washed in nuclei wash buffer (cell lysis buffer without IGEPAL) and then centrifuged again at 500g for 5 minutes 4°C, and the supernatant was discarded. Finally, the pellets were resuspended again in nuclei wash buffer and then flash-frozen in liquid nitrogen before storing in –80°C.

For nuclei extraction from the differentiation culture, the cells were treated with StemPro Accutase (Thermo, #A1110501) for 7min to detach the cells and transfer them into 15mL tube, then incubated in trypsin (Thermo, #25300054) for another 7min to prevent re-clumping. The cells were span down to remove trypsin after inactivation with more media. The pellet was treated with nuclei lysis buffer and the same steps for nuclei extraction protocol were followed.

### Sci-RNA-seq

Single-cell combinatorial-indexing RNA-sequencing (sci-RNA-seq) protocol is described previously (Cao *et al*., 2019). sci-RNA-seq relies on the following steps, (i) thawed nuclei were permeabilized with 0.2% TritonX-100 (Sigma, #T9284) (in nuclei wash buffer) for 3 min on ice, and briefly sonicated to reduce nuclei clumping; (ii) nuclei distributed across 96-well plates; (iii) A first molecular index is introduced to the mRNA of cells within each well, with *in situ* reverse transcription (RT) incorporating the unique molecular identifiers (UMIs); (iv) All cells were pooled and redistributed to multiple 96-well plates in limiting numbers (e.g., 10 to 100 per well) and a second molecular index is introduced by hairpin ligation;(v) Second strand synthesis, tagmentation, purification and indexed PCR; (vi) Library purification and sequencing is performed.

All libraries were sequenced on one NovaSeq platform (Illumina). Base calls, downstream sequence processing and single-cell digital-expression matrix generation steps were similar to what was described in sci-RNA-seq3 paper (Cao *et al*., 2019). STAR (Dobin et al., 2013) v.2.5.2b54 aligner used with default settings and gene annotations (GRCh38-primary-assembly, gencode.v27). Uniquely mapping reads were extracted, and duplicates were removed using the UMI sequence, reverse transcription index, hairpin ligation adaptor index and read 2 end-coordinate (that is, reads with identical UMI, reverse transcription index, ligation adaptor index and tagmentation site were considered duplicates).

### Data Analysis

All low-quality reads were removed from the data (including jaws, toothgerms and salivary glands samples from all time points) by setting UMI cutoff to greater than 200 and removing all mitochondrial reads (QC table: Figure S1F). Following Monocle3 workflow (Cao *et al*., 2019; Qiu et al., 2017; Trapnell *et al*., 2014), data underwent normalization by size factor, preprocessing, dimension reduction (UMAP algorithm (McInnes et al., 2018)), and unsupervised graph-based clustering analysis (Leiden Algorithm (Levine et al., 2015; Traag et al., 2019)). Certain clusters from the initial analysis were selected for further sub-clustering, and the previous analysis repeated. Pseudotime analysis also was done with Monocle3 following the default workflow, which include learning the graph, ordering the cells, and plotting the trajectory over UMAP. Mutual nearest neighbors (MNNs) algorithm (Haghverdi et al., 2018) was used for batch effect correction only between day10 differentiation sample and day16 sample. PanglaoDB (Franzén *et al*., 2019), a curated single-cell gene expression database was utilized to explore the consensus of cell type markers used across publicly available single-cell datasets.

### Top marker genes

Each dataset or subset was analyzed with monocle’s top_maker function to find potential marker genes. All non-protein coding genes, ribosomal and mitochondrial genes were excluded from the input genes, and only the top 100 genes sorter by marker score were included in the results.

### Heatmap and GO-terms enrichment

ComplexHeatmap package (Gu et al., 2016) was used to generate custom heatmaps that integrate GO-terms per clusters. ViSEAGO package (Brionne *et al*., 2019) used to generate the GO-terms, and simplifyEnrichment package (Gu and Hübschmann, 2021) used to extract keywords from the top 100 GO-terms (by p value) per cluster. The top 50 marker genes in each cluster were utilized as the input for ViSEAGO. The keywords generated by simplifyEnrichment, were filtered to eliminate redundant and irrelevant words, and only the very top words are displayed on the heatmap.

### Pseudotime analysis

Pseudotime analysis was done using monocle3 and the density outline of each time point were overlayed on the UMAP graph, to give a better indication of the temporal presence of each cluster.

### Top pathway analysis

We analyzed the stages of ameloblast development as identified in (Figure 3D). OEE and CL were combined as one OEE cluster to increase the statistical power (Figure 5A). To analyze those stages in a thorough and reproducible manner, we compiled a comprehensive analysis pipeline that evaluate pathway activity based on ligand receptor interaction and downstream activity. The workflow for our analysis is shown in (Figure S5A). The first step in the analysis is selecting the appropriate input for each stage of the differentiation to be analyzed. At each stage, we consider the progenitor cells and the target cell type to be differentiated into, as well as all the support cell types that are present in the same stage and that are likely to send the signals. The second step is to analyze all the potential ligand-receptor interactions between the selected cell types, but only focus on in-coming interactions toward the progenitor cells of interest. For this part of the analysis, we used a software, *talklr* package (Wang, 2020), which uses an information-theoretic approach to identify and rank ligand-receptor interactions with high cell type-specificity. We further filtered *talklr* output by selecting those ligand-receptor pairs that fall within the major signaling pathway of interest (TGFβ, BMP, GDF, GDNF, NODAL, ACTIVIN, WNT, ncWNT, EGF, NRG, FGF, PDGF, VEGF, IGF, INSULIN, HH, EDA, NGF, NT, FLT3, HGF, ROBO, NOTCH, NRXN, OCLN). The third step of the workflow is to obtain the differentially expressed genes (DEGs) between the progenitor cells of interest and their differentiated cell type. This set of genes can be used to evaluate the downstream activity and can be linked to specific ligand-receptor pairs. We used DEsingle package (Miao *et al*., 2018) with FDR threshold set to 0.1 to obtain DEGs. The top marker genes for the progenitor cells were also excluded from DEGs in this analysis, to ensure more weight is given to the differentiated cell type. The fourth step is to generate a multilayer network that models the upstream interactions (obtained from step #2) and the downstream interactions that includes transcription factors (TF) and their target genes (DEGs obtained from step #3). We used the R package scMLnet (Cheng *et al*., 2021) to generate the multilayered network interactions that consists of a top layer for ligands, a layer for receptors, a layer for TFs and a layer for TF-targets. The fifth step is to implement a scoring system to evaluate the connectivity of each part of the multilayered network obtained from previous step, to determine which path is more probably active. We started by assigning fold-change values (obtained in step #3) to target genes at the lowest level. At next level, the TF layer, we assigned the mean values of all the connected TF-targets to each TF. Normalization of the scores to the interaction database depth is done after each step, to ensure the scores remain comparable with each category of interactions. At the receptor layer, we calculated the sum of the values of all the connected TFs to each receptor. At the ligand layer, we calculated the sum of the values of all the connected receptors to each ligand. And finally, all ligands that fall within the same pathway family are aggregated together. The sixth step of our pipeline is to rank pathways based on the percentage of activity compared to the overall combined activity scores of all pathways evaluated in this analysis. The results indicate the most active pathways or the most active ligands that are key drivers of the differentiation at a specific stage of development (Figure 5A).

### Differential expression

DEsingle package (Miao *et al*., 2018) was used to calculate differential expression (DE) between clusters. DEsingle was designed for single-cell RNA sequencing, and it employs Zero-Inflated Negative Binomial model to estimate the proportion of real and dropout zeros. Our cutoff for DE genes were set to include genes with False Discovery Rate (FDR) < 0.1 and more than twofold change.

### Multilayer network analysis

To generate a multilayer network that models the upstream interactions and the downstream interactions that includes transcription factors (TF) and their target genes, we used the R package scMLnet (Cheng *et al*., 2021). A custom wrapper code was developed to integrate *talklr* and DEsingle results with scMLnet.

### Signaling interaction

In our study, we used *talklr* package (Wang, 2020) to identify ligand-receptor interaction changes between two adjacent tooth developmental stages. *talklr* uses an information-theoretic approach to identify ligand-receptor interactions with high cell type-specificity. Ligand-receptor interaction score is defined as Li*Rj, the product of expression levels for the ligand in cell type i and the receptor in cell type j. We normalize interaction scores by dividing Li*Rj with the sum of interaction scores across all n2 cell-cell interactions. *talklr* uses the Kullback-Leibler divergence to quantify how much the observed interaction score distribution differs from the reference distribution. The reference distribution is the equi-probable distribution where every possible interaction has probability, when the aim is to identify cell type-specific ligand-receptor pairs in a single condition. Compared to existing methods such as cellPhoneDB (Efremova et al., 2020) or singleCellSignalR (Cabello-Aguilar et al., 2020) the unique strength of *talklr* is that it can automatically uncover changes in ligand-receptor re-wiring between two conditions (e.g. different time points, disease vs. normal), where the reference distribution is the observed interaction scores in the baseline condition. The parameters we used were 0.001 for expression threshold, which was determined by calculating the level of expression of the 20th quantile of the aggregated clusters, and 1e-06 for the pseudo-count value which was determined by the minimum averaged expression value in the set. We considered the interactions among the top 100 ligand-receptor pairs returned by *talklr*, and we further prioritized them by selecting those that are known to be from physically proximal cell types.

### Datasets projection analysis

Seurat 4.0 package (Hao *et al*., 2021) was used to project the *invitro* differentiation sample into the UMAP space of fetal ameloblasts sample. The dataset in monocle object format that contains the precomputed PCA and UMAP was converted into Seurat object. The projection was done with the default parameters. Graph-based clustering was performed on the projected data by calculating the nearest neighbor cluster center of the fetal sample. Package ‘networkD3’ (Allaire et al., 2017) was used to create the river plot showing the proportions of the classified cells.

### Datasets integration analysis

LIGER package (Welch *et al*., 2019) was used to integrate the fetal dental epithelium lineage dataset with the differentiation datasets to facilitate the cell type label transfer between the sets. The following integration parameters were used: k = 25, lambda = 10, and these settings were determined by utilizing the built-in function that suggest the best values that suit our datasets. For the river plot generation, the minimum fraction of the branching streams was set to 0.25, and the minimum number of cells set to 50. Clusters that have no out-or ingoing connection were eliminated from the graph for clarity.

### RNA Fluorescence *in situ* Hybridization (FISH) and analysis

A 12-probe RNAScope HiPlex assay (Advanced Cell Diagnostics, Inc.) including probes against 13 transcripts differentially expressed between cell type clusters in mesenchyme-and epithelial-derived lineages were selected to distinguish cell populations: *VWDE, SALL1, FGF4, IGFBP5, FGF10, PRRX1, FBN2, ENAM, PCDH7, SOX5, KRT5,* and either *DSPP or LGR6*. Fresh frozen tissue sections from d80 and d117 were assayed according to the manufacturer’s protocol. Briefly, the fresh-frozen tissue sections were fixed using 4% paraformaldehyde in 1X PBS, dehydrated, and treated with the Protease IV kit component. The first four probes were imaged after completing the manufacturer’s specified hybridization steps, counterstaining, and coverslipping. Images of tissue sections were obtained using an Nikon Ti2 with an Aura light engine (Lumencor, Beaverton, OR), and BrightLine Sedat filter set optimized for DAPI, FITC, TRITC, Cy5 & Cy7 (Semrock, Rochester, NY: LED-DA/FI/TR/Cy5/Cy7-5X5M-A-000) or a Yokogawa CSU-X1 spinning disk confocal microscope (Yokogawa Corporation, Sugar Land, TX) with a Celesta light engine (Lumencor), ORCA-Fusion scientific CMOS camera (Hamamatsu Corp, Bridgewater, NJ), and a HS-625 high speed emission filter wheel (Finger Lakes Instrumentation, Lima, NY). Coverslips were removed, the first four fluorophores were cleaved, and the process was repeated for probes 5-8 and then probes 9-12 (File S4). Images were analyzed using Fiji (ImageJ2 v2.3.0) and QuPath (v0.3.0) quantitative pathology and bioimage analysis freeware (Bankhead et al., 2017). Briefly, The DAPI channel images for imaging rounds two and three were aligned to the DAPI image for imaging round one using the BigDataViewer > BigWarp plugin in Fiji. Matching reference points were identified across the DAPI images and the resultant landmark tables were used in a custom .groovy script (File S5) to align the FITC, Cy3, Cy5, and Cy7 images from the three rounds of imaging. Images were uniformly background corrected and scaled as indicated in File S4. Cellular segmentation was performed in QuPath and positive signal foci and clusters were identified as subcellular detections. Parameters were set to allow for detection of foci while avoiding false positive detection events using positive and negative control images. From QuPath, the coordinates and the number of spots estimated (sum of individual puncta and estimated number of transcripts for clustered signal) for each segmented cell were processed using custom R scripts to map cell locations and expression levels. Out of the transcripts assayed by RNAScope, probe set criteria (File S3) used to identify a given cell population in RNAScope data was selected based on differential expression across the cell types identified in the sci-RNA-seq data at corresponding time points (Figure S3C). Cells matching expression criteria for a cluster’s probe set were designated by cluster color and mapped spatially.

### *In vitro* differentiation

Briefly, hiPSCs (WTC-11 human induced pluripotent stem cells) (Coriell, #GM25256) were seeded on 12-well plates coated with growth factor-reduced Matrigel (Corning, #356231) and cultured in mTeSR1 stem cell medium (StemCell Technologies, #85850) until cells reach confluency with medium changes daily. On the first day of differentiation (deemed Day 0), stem cell media is replaced with ameloblast base media consisted of either EpiCult-C media (StemCell Technologies, #05630) or RPMI 1640 Medium (Thermo, #11875093) mixed with EpiLife (Thermo, #MEPI500CA) at 1:1 ratio, supplemented with 0.1x supplement S7 (Thermo, #S0175), 0.1uM β-mercaptoethanol (BME) (Sigma, #M7522) and 400um smoothened agonist (SAG) (Selleckchem, # S7779). At day 3 of differentiation 150pM of bone morphogenic protein-4 (BMP4) (rndsystems, #314-BP-010) is continuously added daily till day 7. At day 8, the base media is supplemented with 1uM of BMP-I inhibitor (LDN-193189) (Tocris, # 6053), 5uM of GSK3-Inhibitor (CHIR99021) (Selleckchem, # 4423), 500pM epidermal growth factor (EGF) (rndsystems, #236-EG) and 3.5μM of Neurotrophin-4 (NT4) (rndsystems, #268-N4). The cultures were then harvested at day 10 at an oral epithelium stage, or extended to day 16 by adding 300pM BMP4, and 800nM transforming growth factor beta 1(TGFβ1) (rndsystems, #7754-BH) for the early ameloblast stage at day16. For testing FGFR signaling requirement for the maturation process we added 50nM purified FGFR-mb (see below) to the media at day 14 and harvested the samples at day 16 of the differentiation.

### De novo FGFR-Miniprotein expression

The gene encoding the designed FGFR-mb protein sequence was synthesized and cloned into modified pET-29b(+) *E. coli* plasmid expression vectors (GenScript, N-terminal 8-His tag followed by a TEV cleavage site). The sequence of the N-terminal tag is MSHHHHHHHHSENLYFQSGGG, which is followed immediately by the sequence of the designed protein. Plasmids were transformed into chemically competent *E. coli* Lemo21 cells (NEB). The protein expression was performed using Studier autoinduction medium supplemented with antibiotic, and cultures were grown overnight. Then, IPTG was added to a final concentration of 500□mM and the cells were grown overnight at 22□°C for expression. The cells were collected by spinning at 4,000*g* for 10□min and then resuspended in lysis buffer (300□mM NaCl, 30□mM Tris-HCL (pH□8.0), with 0.25% CHAPS for cell assay samples) with DNase and protease inhibitor tablets. The cells were lysed with a sonicator (Qsonica Sonicators) for 4□min in total (2□min each time, 10□s on, 10□s off) with an amplitude of 80%. The soluble fraction was clarified by centrifugation at 20,000*g* for 30□min. The soluble fraction was purified by immobilized metal affinity chromatography (Qiagen) followed by FPLC SEC (Superdex 75 10/300 GL, GE Healthcare). The protein samples were characterized by SDS–PAGE, and purity was greater than 95%. Protein concentrations were determined by absorbance at 280□nm measured with a NanoDrop spectrophotometer (Thermo Scientific) using predicted extinction coefficients.

### RNA extraction and RT-qPCR analysis

RNA was extracted using Trizol (Life Technologies) according to manufacturer’s instructions. RNA samples were treated with Turbo DNase (Thermo Fisher Scientific) and quantified using Nanodrop ND-1000. Reverse transcription was performed using iScript cDNA Synthesis Kit (Bio-Rad). 10□ng of cDNA was used to perform QRT-PCR using SYBR Green (Applied Biosystems) on a 7300 real time PCR system (Applied Biosystems). The PCR conditions were set up as the following: stage 1 as 50□°C for 2□mins, stage 2 as 95□°C for 10mis, 95□°C for 15□sec, 60□°C for 1□min (40 Cycles). ß-actin was used as an endogenous control. The primer sequences used in this work are available in File S6.

### Development of Ameloblast Organoid

The day16 differentiated iAM cells were trypsinized using TrypLE (Thermo Scientific) and re-plated in in 24-well ultra-low attachment plate (Corning, #4441) containing an ameloblast base medium with 10 μM ROCKi (Y-27632, Selleckchem, #S1049). The organoid cultures were maintained at 37°C in 5% CO2, and the medium was changed every 3-days until further analysis.

### Co-culture protocol for ameloblast and odontoblast organoid

The day16 differentiated iAM cells were cultured in ultra-low attachment 12-well plate for a week in ameloblast base medium. The odontogenic organoids were made in a similar manner in a separate plate by culturing DPSCs

(isolated from primary molar sample of young patient (Macrin et al., 2019)) in odontogenic differentiation medium containing DMEM (Gibco, #11995073) ascorbic acid (Sigma, #A8960), β-Glycerophosphate (Sigma, #35675), and dexamethasone (Sigma, #D2915), 10% FBS (Gibco, #10437028) and 1% Penicillin/Streptomycin (Gibco, #15140122). The two types of organoids were co-cultured in the same wells for two weeks, supplemented with a 1:1 mixture of both odontogenic and ameloblasts base media at 37°C in 5% CO2. The co-culture was sampled later for further analysis.

### Co-culture protocol for monolayer

The DPSCs were plated as monolayer mixed in 25% (v/v) of Matrigel (Corning, #356231) diluted in odontogenic media in a glass-bottomed 96-well plate (Corning, #3603). The following day, iAM cells suspended in the ameloblast base medium and 10 μM ROCKi (Y-27632, Selleckchem, #S1049) were added on top of the DPSCs monolayer and then incubated for 24 hours at 37°C in 5% CO2. The formed organoids were supplemented with fresh media (1:1 mixture ameloblast and odontogenic media) containing Calcein solution (Sigma, #C0875) (1uM, 1:1000) on every three consecutive days. The co-culture was sampled on the 14th day for further analysis.

### Cryosectioning and Immunostaining for the organoids

The organoids were imbedded in OCT compound (Tissue-Tek, # 4583) and slowly frozen on a metal block chilled on dry ice. Frozen organoids were cut using Cryostat (Leica CM1850) to create 10μm slices and fixed on glass slides (Fiserbrand, #12-55015) for staining. The organoid sections were fixed in 4% paraformaldehyde (PFA) for 10-15min at RT and later washed thrice with 1X PBS for 5 min each. Slides were then immersed in 0.5% TritonX 100 at RT for 5 minutes to facilitate permeabilization. Later blocking was done for 1hour at RT in a humidified chamber with a blocking buffer consisting of 0.1% Triton X-100 and 5% Bovine Serum Albumin (VWR). The organoids were incubated in primary antibodies (File S6) overnight at 4□ in a humidified chamber. After 3x5 minute washes in PBS in a coplin jar, the slides were transferred to a humidified chamber with secondary antibodies. Secondary antibodies and Phalloidin (File S6) were applied for 1hour at RT in the same blocking agent, followed by rinsing the slides with PBS 3x5min in coplin jar. The slides were incubated in autofluorescence quenching solution (Vector Labs, #SP-8400) for 5 min at RT under dark conditions and rinsed 1x with PBS. DAPI (Thermo Fisher) was applied for 10 minutes at room temperature in PBS. Slides were then rinsed with PBS for 10 minutes in a coplin jar. Slides were then mounted with Vectashield (Vector Labs) and stored at 4□ for imaging.

### Wholemount immunostaining analysis

The organoids were collected in a 2ml tube after two weeks and washed thoroughly with 1x PBS before fixation. The organoids were fixed in 4% paraformaldehyde (PFA) for 10-15min at RT on a rocker. Later the fixed organoids were washed thrice with 1X PBS for 5 min each. The organoids were then immersed in 0.5% TritonX 100 at RT on a rocker for 5 minutes. Later blocking was done for 1hour at RT on a rocker with a blocking buffer consisting of 0.1% Triton X-100 and 5% goat serum (VWR). The organoids were incubated overnight in the primary antibodies (File S6) at 4°C on a rocker. After 5-minute washes in PBS for thrice in a coplin jar, the organoids were incubated with secondary antibodies (File S6) for an hour at RT on a rocker. The primary and the secondary antibodies were prepared in the blocking agent consisting of 0.1% Triton X-100 and 3% goat serum (VWR). Followed by washing the organoids with PBS 3x5min on a rocker. The organoids were incubated in autofluorescence quenching solution (Vector Labs, #SP-8400) for 5 min at RT under dark conditions on a rocker and rinsed 1x with PBS. Incubate the organoids in 200 mL of PBS containing DAPI (Thermo Fisher) for 10 min. The organoids were then rinsed with PBS, mounted with Vectashield (Vector Labs, # H-1700), and stored at 4□ for imaging.

### Injection of iPSC-derived ameloblast-like cells into mouse muscles

hiPSCs (WTC11) were allowed to undergo differentiation for the pre-ameloblast stage at day16 using the following basal supplements mentioned above cultured in Matrigel. 1 × 10^6^ iAM cells were resuspended in Matrigel supplemented with a cocktail of prosurvival factors (Laflamme et al., 2007) and injected into the femoral muscle of SCID-Beige mice (Charles River, Wilmington, MA). Mice were kept under BioSafety containment

Level 2. Mice were sacrificed and femoral muscles were harvested after 2 months and were dissected at the site of injection (left leg muscle) to perform further analysis. Experiment was performed in compliance with ethical regulations, IACUC protocol #4152-01. After dissection, left leg muscles were embedded in embedding cryo-mold (Polysciences, #18986-1) with minimum amount of Tissue-Tek O.C.T. compound (Sakura, catalog number: 4583) to cover the muscle region. The embedded tissue was then snap-frozen by placing on a cold-resistant beaker of 2-methylbutane solution (EMD. #MX0760-1) into a slurry of liquid nitrogen for 5-mins, which allows fast cooling to -80 °C. The snap-frozen samples are then placed in a -80 °C freezer for storage. The cryostat and blade are both pre-chilled to -20°C before cryo-sectioning. 10 μm-thick sections were made on pre-chilled Superfrost Plus microscope slides (Fisherbrand, #12-550-15) and then store in a -80 °C.

### Calcification assays: Von Kossa and Alizarin Red Staining

Identification of mineralization was performed on tissue sections stained with Von Kossa and Alizarin Red S. Frozen leg muscle sections (10μm) were fixed with 4% paraformaldehyde (EMS, #15710) in H_2_O at room temperature for 12min. Rinse the section with deionized distilled water thrice for 5min each. Sections were incubated in with 5% silver nitrate solution (SIGMA-ALDRICH #209139) placed under ultraviolet light for 1 hour. The section was rinsed with several changes of deionized distilled water for 5min each and later incubated in 5% Sodium Thiosulfate solution (SIGMA-ALDRICH #217263) for 5 minute to remove un-reacted silver. Similarly, sections were stained with 2% Alizarin red S solution (pH4.2) (Sigma, #A5533) for 1 hour in the dark. The slides were thoroughly with deionized distilled water for 5min each followed by counterstaining the sections with nuclear fast red stain (EMS, # 26078-05) for 5 minutes. Rinsed in deionized distilled water briefly for 5mins each the slides were successfully transferred into coplin jars to perform dehydration step through graded alcohol and clear the slides in CitriSolv solution (Decon, #1601). Slides were then mounted with Vectashield (Vector Labs, #H-1400-10) and stored at room temperature for imaging.

### Cryosectioning of fetal samples

Jaw tissues were fixed with 4% PFA overnight at 4°C followed by 30% sucrose (Sigma, #RDD023) treatment until the tissue sank to the bottom of the tube. The tissue is then imbedded in OCT compound (Tissue-Tek, # 4583) and slowly frozen on a metal block chilled on dry ice. Frozen samples were cut using Cryostat (Leica CM1850) to create 10μm slices of tissue and fixed on glass slides (Fiserbrand, #12-55015) for staining.

### Immunofluorescence staining and Confocal Imaging

Toothgerms embedded in O.C.T. were cryosectioned to 10-micron thick sections. The slides were stored at -80□ after cryosectioning and warmed at room temperature prior to staining. Tissues were fixed in 4% paraformaldehyde (PFA) then immersed in 1X PBS for 3x5 minute washes. Antigen retrieval was performed using 10X Citrate Buffer (Sigma-Aldrich) in a capped coplin jar microwaved for ∼45 seconds followed by 15-minutes incubation in microwave. Slides were then allowed to be washed in PBS at room temperature for 7 minutes. Slides were blocked for 90 minutes at room temperature in a humidified chamber with a blocking buffer consisting of 0.1% Triton X-100 and 5% Bovine Serum Albumin (VWR). All the antibodies used in this study and their concentrations are listed in File S6. The primary antibodies were incubated overnight at 4□ in a humidified chamber. After 3x5 minute washes in PBS in a coplin jar, the slides were transferred to a humidified chamber with secondary antibodies. Secondary antibodies were applied for 75 minutes at room temperature in the same blocking agent. Slides were then rinsed with PBS 4x10 minute washes in a coplin jar. DAPI (Thermo Fisher) was applied for 10 minutes at room temperature in PBS. Slides were then rinsed with PBS for 10 minutes in a coplin jar. Slides were then mounted with Vectashield (Vector Labs) and stored at 4□ for imaging. Confocal Imaging was done on a Leica TCS-SPE Confocal microscope using a 40x objective and Leica Software. Images were processed with Fiji software distribution of ImageJ v2.3.0 (Schindelin et al., 2012; Schindelin et al., 2015). NIS-Elements (RRID:SCR_014329) was used for 3D reconstruction.

## Data availability

The data generated in this study can be downloaded in raw and processed forms from the NCBI Gene Expression Omnibus under accession number (GSE184749). Upon publication, raw RNAScope data will be made publicly available on <u>dryad.org</u> (Dryad research data repository).

## Code availability

The custom R codes used to generate some of the results in this paper are available in https://github.com/Ruohola-Baker-lab/Tooth_sciRNAseq.

## Supporting information

File S1_marker_genes

File S2_Meta-analytic_estimators

File S3_RNAscope_logic_tables

File S4_QuPath_Settings

File S5_align-to-R1-batched.groovy

File S6_antibodies_and_primers

## Acknowledgment

We thank the Ruohola-Baker lab members for helpful discussions during the course of this work. We thank Chris Cavanaugh, Fatima Al-Shimmary, Christopher Kelley, Aaron Liu, Zicong Lee, Gwen L. Tilmes and Anoushka Amath for their excellent technical help. We thank Prof. Carol Ware for inspiring discussions and guidance. We thank Kimberly A. Aldinger, Ian G. Phelps, Jennifer C. Dempsey, Kevin Lee and Lucy Cort from BDRL for processing and collecting the samples from the donors and troubleshooting the nuclei isolation. We thank the BBI for the technical support and overall guidance in the design of the sequencing project. This work is supported by grants from the National Institutes of Health 1P01GM081619, R01GM097372, R01GM97372-03S1 and R01GM083867 and the NHLBI Progenitor Cell Biology Consortium (U01HL099997; UO1HL099993) for HRB. This work was supported by the BBI grants for JM, YW and HRB. This work was also supported by Dr. Douglass L. Mourell Research Fund for HRB, HZ, AA, and YTZ. The Birth Defects Research Laboratory was supported by NIH award number 5R24HD000836, to IAG, from the Eunice Kennedy Shriver National Institute of Child Health and Human Development. AA was supported by Imam Abdulrahman bin Faisal University, and Saudi Arabian Cultural Mission (SACM) to the USA. SHD, YTZ and DE were supported by National Institutes of Health T90DE021984, and SHD was also supported by ARCS. MCR and work conducted in the ISCRM Genomics Core were supported by a generous gift from the John H. Tietze Foundation.

---

**Figure S1:**
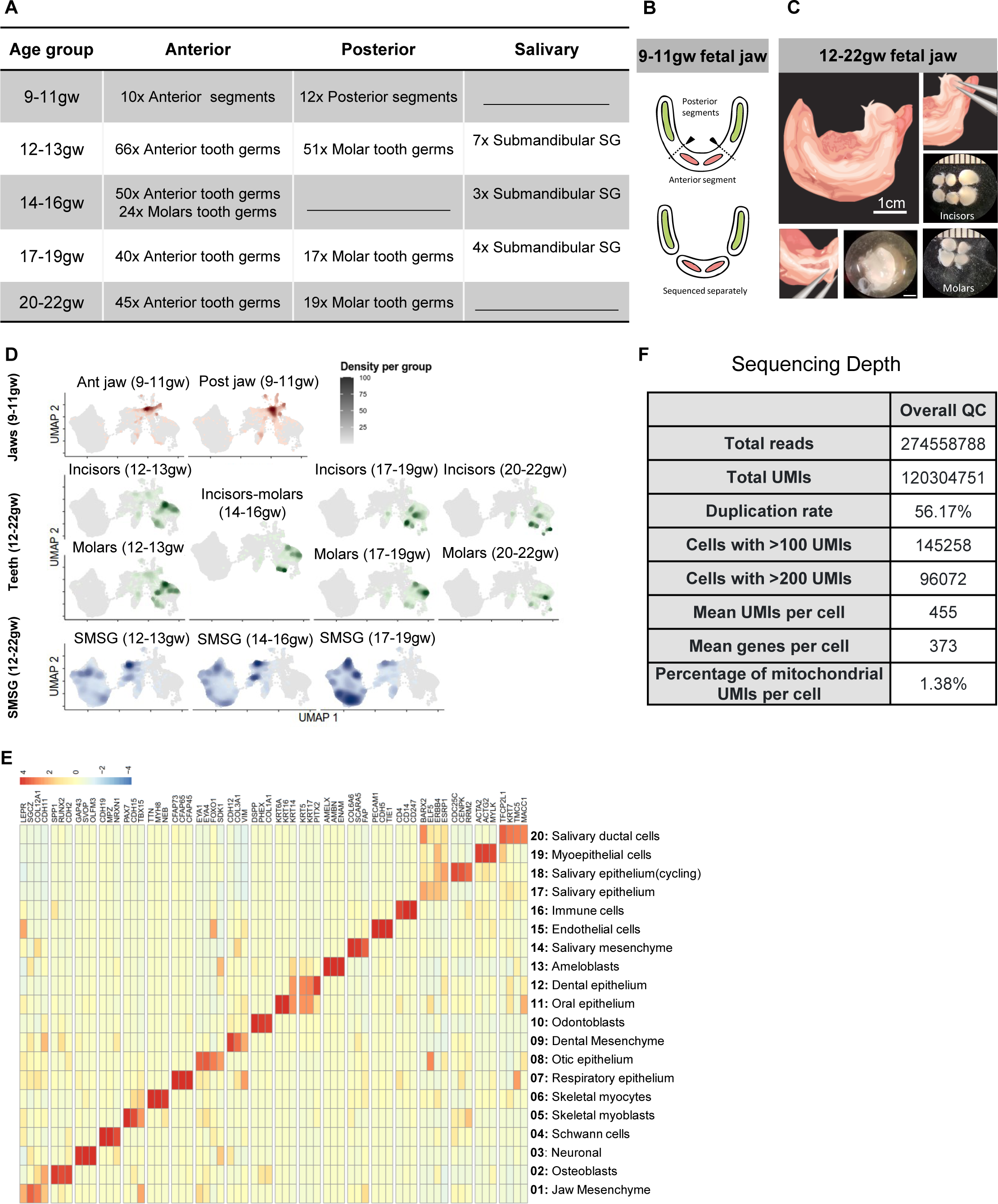
A single-cell atlas of the developing human fetal jaws, teeth, and salivary glands tissues via sci-RNA-seq. (related to Figure 1) (A) Depending on the tissue size at a given age, between 3 and 50 of each tissue sample were collected, pooled into 12 samples, and sent for sci-RNA-seq. (B) At 9-11w, dissecting individual toothgerms or salivary glands in the bud stage was not feasible due to the large number of cells required to perform sci-RNA-seq protocol. Instead, jaws were separated into two segments of posterior jaw, containing jaw tissue distal of the canines (B, red boxes), and one segment of anterior jaw spanning from canine tooth to canine tooth region (B, blue box). At 12 weeks gestation and beyond, individual toothgerms and submandibular salivary glands could be identified and isolated to be sequenced separately (C). Sequenced data was clustered, and the resulting plot revealed that each major tissue type occupied a specific region of the plot, with some shared support tissues localized in the center (D). Clusters were identifiable by expression of known markers for each tissue type in heatmap (E). QC table for all sequenced data (F).

**Figure S2:**
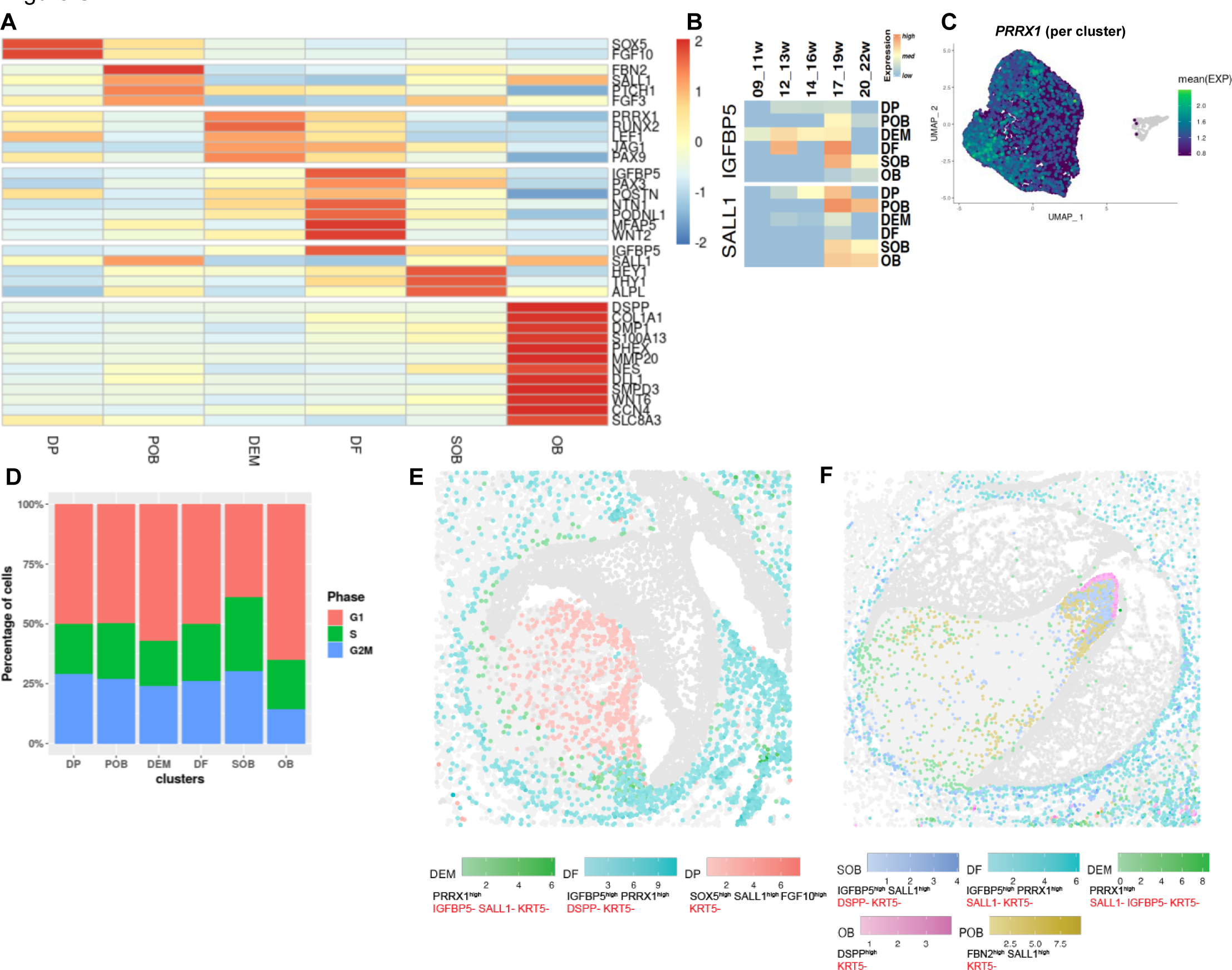
Expression of known marker genes for dental mesenchyme derived cell types. (related to Figure 2) (A) Heatmap of expression over time of dental follicle marker *IGFBP5* and subodontoblast markers *SALL1* (B). Gene plot of shared DP and DEM progenitor marker *PRRX1* (C). Cell cycle scoring of dental mesenchyme derived cell types (D). Mappings for dental mesenchyme-derived cell types at 80d replicate (E) and 117d replicate (F) identified by analysis of RNAScope images which show mapping for SOB, DF, DEM, OB, and POB cell types.

**Figure S3:**
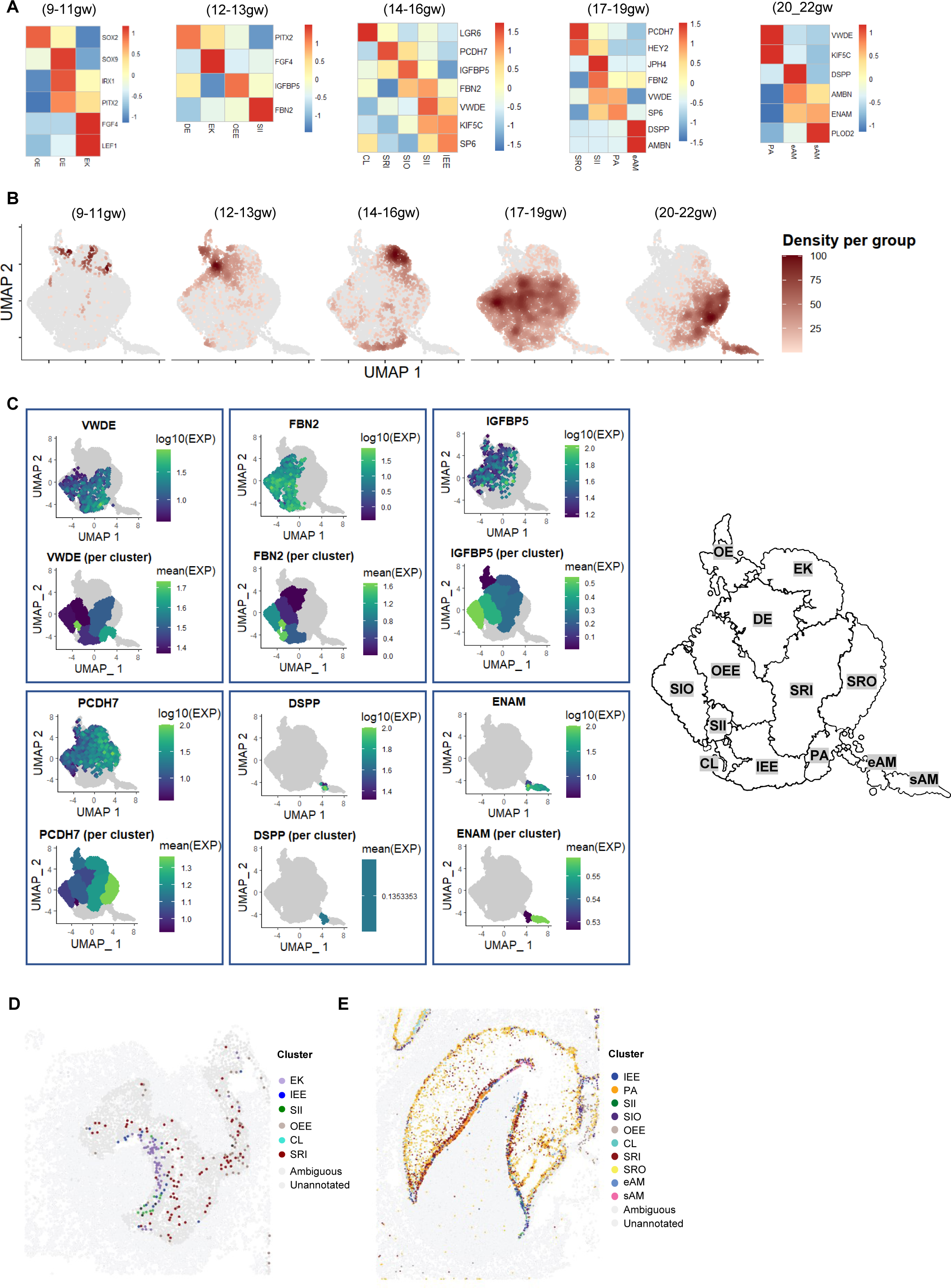
Ameloblast developmental trajectory. (related to Figure 3) (A) Expression of known markers at stages of dental epithelial lineage align with previously identified markers in each tissue and appear at expected developmental timepoints (A). Density of cells plotted by age demonstrates that clusters enriched with more cells at a given timepoint (B). Gene plots and mean expression per cluster summary plots in UMAP space (C) generated for the markers used to infer the logic table (File S3) which is used in the RNAScope mapping. The threshold expression per cluster was set to 25% of maximum expression per gene. Light green clusters considered as high expressing, dark blue as low expressing, and gray as low or no expression. Mappings for dental epithelium-derived cell types in a stage matched (80d) replicate sample (D) identified by analysis of RNAScope images showing the mappings for EK, OEE, IEE, CL, SII, and SRI cell types in composite, and at (117d) replicate sample (E) mappings for IEE, PA, SII, SIO, OEE, CL, SRI, SRO, eAM, and sAM cell types at 117d shown in composite.

**Figure S4:**
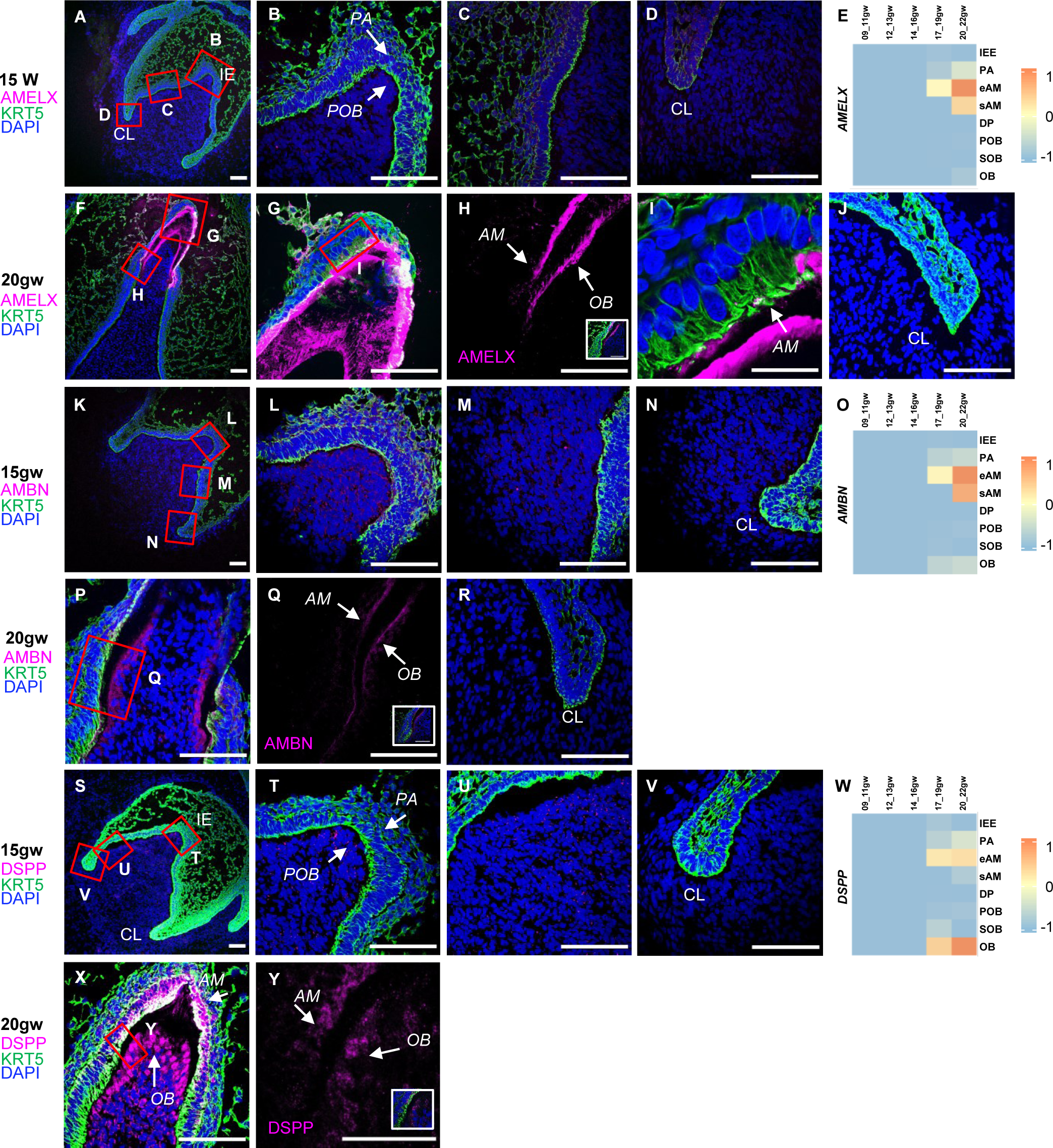
Spatial Expression of Odontoblast and Ameloblast Markers Differs Markedly from Early to Late Toothgerm Development. (related to Figure 2 and 3) Ameloblast markers amelogenin (AMELX) and ameloblastin expression begins in the ameloblast after the early bell stage (A-J, K-R). Similarly, odontoblast marker dentin sialo phosphoprotein (DSPP) begins in the odontoblast after the early bell stage (S-Y). Heatmaps of expression over time of AMELX (E), AMBN (O), and DSPP (W). AMELX, AMBN and DSPP show mirrored expression patterns in ameloblasts and odontoblasts at late bell stage (H,Q,Y). Abbreviations: preodontoblast (POB), odontoblast (OB), preameloblast (PA) ameloblast (AM), incisal edge (IE), cervical loop (CL). Scale bars: 50μm.

**Figure S5:**
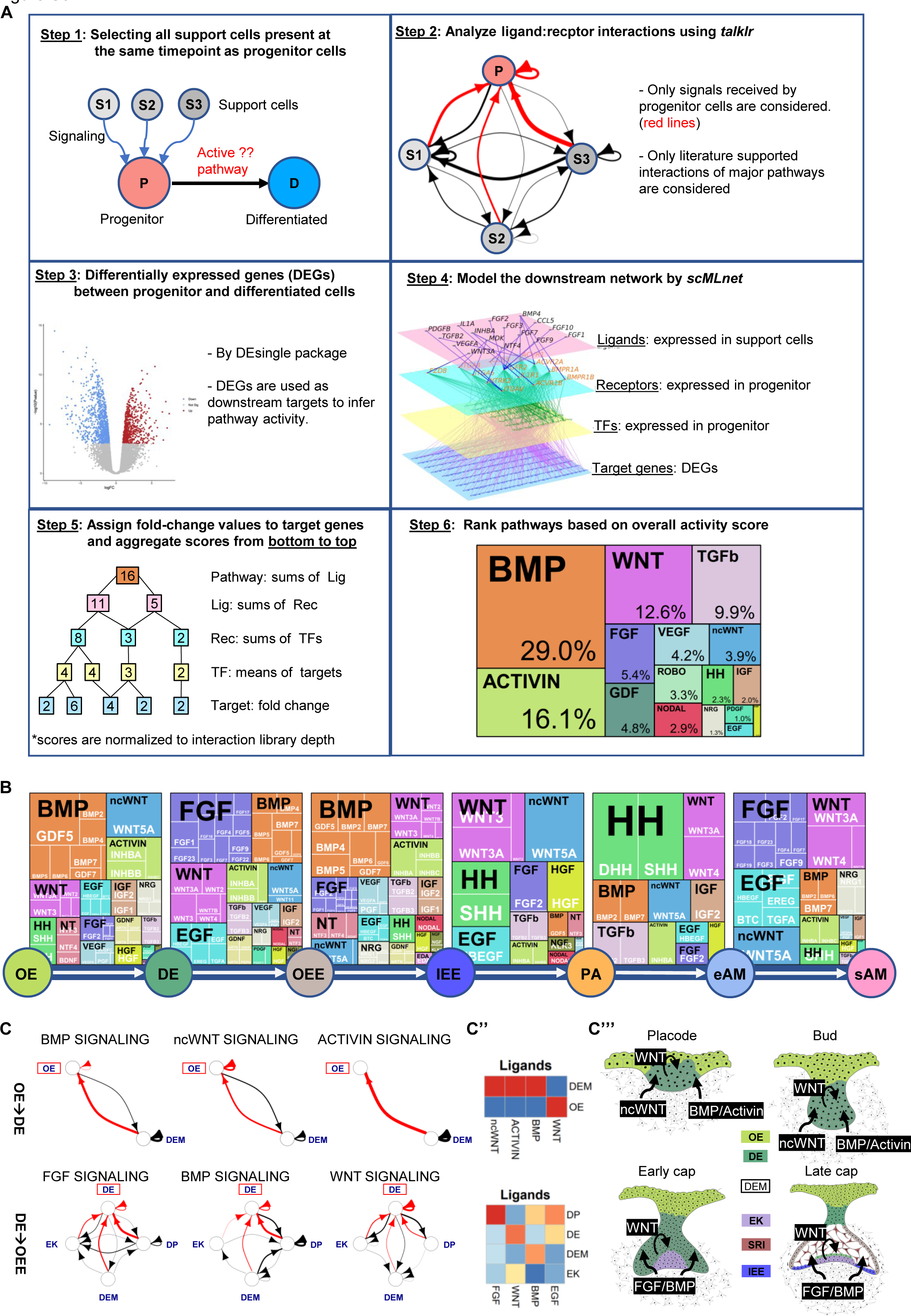
Pathway analysis in ameloblast trajectory. (related to Figure 4) (A) We developed a combined computations workflow to identify critical pathways at different developmental steps, represented by the schematic in (A), yielding a ranking and contribution to ameloblast differentiation for several major pathways (B) over several developmental stages. Using *talklr* ligand-receptor analysis, we identified the sources of the outgoing signals at early stages of ameloblast development (C-C’’’). The top three pathways per stage are indicated in ligand:receptor interaction graphs (C). The thickness of the arrows indicates the number of unique possible interactions between clusters. The arrowheads indicate the receiver cells that express the receptors. Red arrows highlighting the interactions received by the cells of interest at each stage. Heatmaps generated by first aggregating ligands gene expression for each cell, and then the average values are calculated per cluster (C′′). Analyses demonstrate that at the placode stage, BMP and Activin signals come from the mesenchyme (DEM) to the epithelium while at the bud stage the oral and dental epithelium itself secrete WNT signals. In the early cap stages, WNT signals come from the dental epithelium while the mesenchyme secretes FGF and BMP signals toward the dental epithelium. At the late cap stage, following the development of the inner stellate reticulum (SRI), WNT signals switch to non-canonical WNT signaling from the SRI to the enamel knot (C′′′).

**Figure S6:**
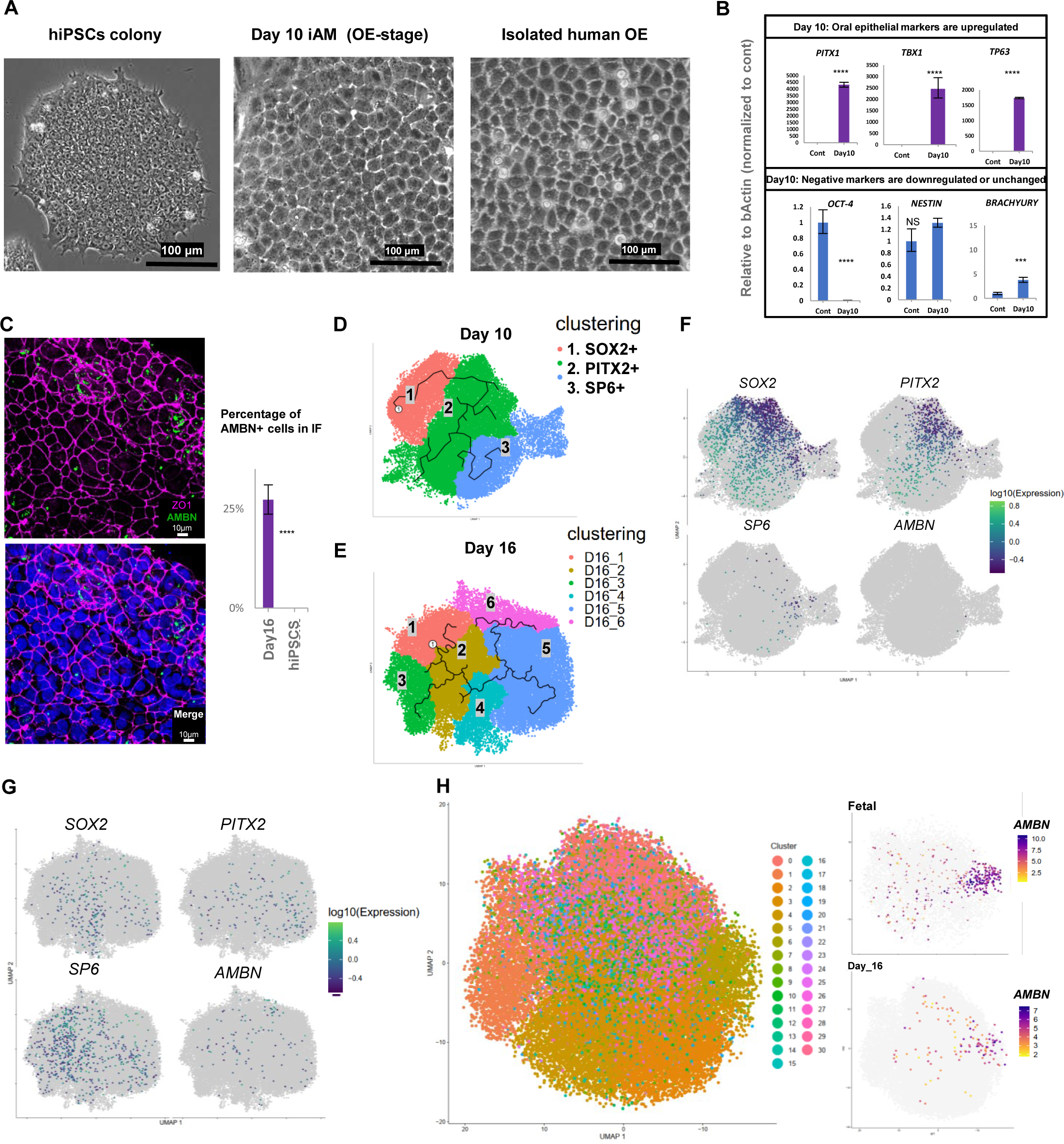
Differentiation of human induced pluripotent stem cells (hiPSCs) into pre-ameloblasts. (related to Figure 5) (A) Brightfield images of hiPSCs, day 10 of *in vitro* differentiation, and isolated fetal oral epithelium after culturing for seven days (A) show that oral epithelium differentiated from iPSCs exhibit the same morphological characteristics as culture human oral epithelium. Quantitative PCR (B) showed that compared to undifferentiated hiPSCs, differentiated oral epithelium exhibited elevated levels of known oral epithelium markers concomitant with a significant decrease in known pluripotency marker *OCT-4*. Additionally, the neuroepithelial marker *NESTIN*, and the early mesodermal marker *TBXT* (BRACHYURY) are relatively unchanged at day10 of the differentiation, indicative of a relatively lineage-specific differentiation. Successful further differentiation of oral epithelium into ameloblasts was demonstrated by immunofluorescence staining of day 16 (C), showing AMBN expression, and the membrane marker ZO1, with quantification analysis finding approximately 25% of cells positive for AMBN expression. Each study was performed in triplicate, with error bars representing ±SEM. Significance was determined by unpaired Student’s t-test; ***p□<□0.001; ****p□<□0.0001. Day 10 (D) and Day 16 (E) samples were sequenced with sci-RNA-seq. Cells were clustered and analyzed to identify clusters with similar gene expression patterns to known cell types in fetal development. Gene expression density plots for known markers of different phases of ameloblast development show continuity between day 10 and day 16, with early markers SOX2 and *PITX2* being predominantly expressed by day 10 (F) and shifting toward ameloblast-specific markers *AMBN* and *SP6* in day 16 (G). LIGER joint clustering analysis of Day 16 differentiation cells and the *in vivo* human fetal dental epithelium (Figure 3A) derived cells suggests the colocalization of *AMBN* expressing cells from in *vivo* and in *vitro* in cluster 6(H).

**Figure S7:**
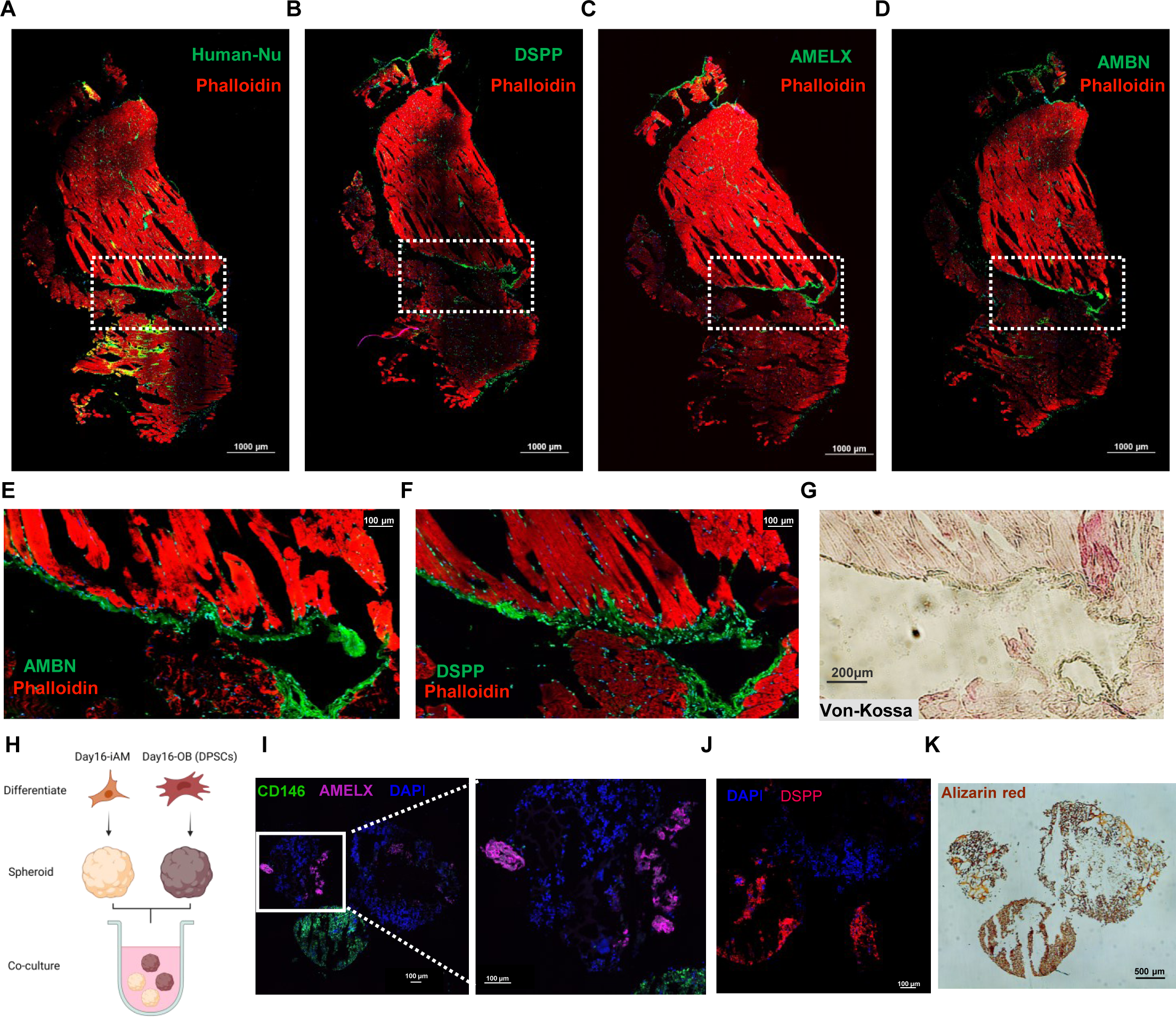
Characterization of iAM and formation of iAM/OB organoids. (related to Figure 6) (A-D) Immunofluorescence staining shows the entire surface area of the stained muscle sections in Figures 6B– 6E. The dotted boxes indicate the area of interest magnified in main Figure 6 and in (S7E–S7G) for AMBN and DSPP. Von-Kossa staining for calcification was performed in a subsequent section in (G) showing black/brown staining localized to the injected cell region. (H) A schematic for the coculture experiment between iAM organoids and DPSC organoids in suspension culture where each cell type was formed in separate wells and then combined for 14 days in iAM base media. Organoids were snap frozen, cry-sectioned, and prepared for immunofluorescence. Expression of AMELX was noted in iAM organoids (I) and CD146 and DSPP in DPSC/OB organoids (J). Alizarin red staining in (K) indicates the classification is positive in both organoid types, particularly DSPC/OB, which shows more calcifications.

**File S1**. Marker genes used to identify cell types in Figure.1D; Figure 2A and Figure 3A in separate excel sheets.

**File S2**: Meta-analytic estimators for mesenchymal and epithelial lineage related to Figure 2B and Figure 3B in separate excel sheets

**File S3**: RNAScope_logic_tables.xlsx – Criteria used to locate cell types using RNAScope *in situ* hybridization signals

**File S4**: QuPath_Settings.xlsx – Settings used to quantify RNAScope HiPlex *in situ* hybridization signal using QuPath software

**File S5**: align-to-R1-batched.groovy – Script used to uniformly align the FITC, Cy3, Cy5, and Cy7 images from the three rounds of RNAScope HiPlex *in situ* hybridization imaging in Fiji

**File S6**: Antibody list and primer list

