## Supplementary material for "Human iPSC Derived Enamel Organoid Guided by Single-Cell Atlas of Human Tooth Development": File S5_align-to-R1-batched.groovy: File S5_align-to-R1-batched.groovy

```
#@ File (label="Landmark file directory", style = "directory") landmarksDir
#@ File (label="Image directory", style = "directory") Dir
#@ String  (label = "Round", choices={"R2", "R3"}) roundString
#@ String  (label = "Identifier", value = "") identString
#@ File    (label = "Output directory", style = "directory") dstFile
#@ String (label="Transform type", choices={"Thin Plate Spline", "Affine", "Similarity", "Rotation", "Translation" }) transformType
#@ String (label="Interpolation", choices={"Linear", "Nearest Neighbor"}) interpType
#@ Integer (label="Number of threads", min=1, max=64, value=5) nThreads
#@ Boolean (label="Virtual stack?") isVirtual


import java.io.File;
import java.io.IOException;
import bdv.ij.ApplyBigwarpPlugin;
import bdv.viewer.Interpolation;
import bigwarp.landmarks.LandmarkTableModel;
import ij.IJ;
import ij.ImagePlus;


def target() {
	Dir.eachFileRecurse {
		targetname = it.getName()
		if (targetname.contains(identString) && targetname.contains("R1") && targetname.endsWith("1.tif")) {
			landmark(it)
		}
	}
}

def landmark(tar) {
	landmarksDir.eachFileRecurse {
		lmname = it.getName()
		if (lmname.contains(identString) && lmname.contains(roundString)) {
		 moving(tar, it)
		}
	}
}

def moving(tar,lm) { 
	Dir.eachFileRecurse {
		movname = it.getName()
		if (movname.contains(identString) && movname.contains(roundString)) {
			process(it, tar, lm, dstFile)
		}
	}
}


def process(file, tar, lm, dst) {
	println "Processing $file"

	// Opening the image
movingIp = IJ.openImage(file.getAbsolutePath());
targetIp = IJ.openImage(tar.getAbsolutePath());

int nd = 2;
if ( movingIp.getNSlices() > 1 )
	nd = 3;

ltm = new LandmarkTableModel( nd );
try
{
	ltm.load( lm );
} catch ( IOException e )
{
	e.printStackTrace();
	return;
}

Interpolation interp = Interpolation.NLINEAR;
if( interpType.equals( "Nearest Neighbor" ))
	interp = Interpolation.NEARESTNEIGHBOR;

emptyWriteOpts = new ApplyBigwarpPlugin.WriteDestinationOptions( "", "", null, null );
warpedIpList = ApplyBigwarpPlugin.apply(
		movingIp, targetIp, ltm, transformType,
		"Target", "", "Target",
		null, null, null,
		interp, isVirtual, nThreads, true, emptyWriteOpts );

warpedIpList.get(0).show();
transformed = warpedIpList.get(0);
	
	
	println "Saving $file"
	// Saving the result

	saveDir = new File(dst.toPath().toString())
	if (!saveDir.exists()) saveDir.mkdirs()
	saveFile = new File(saveDir, file.getName()) // customize name if needed
	IJ.saveAs(transformed, "Tiff", saveFile.getAbsolutePath());

}

target()
```
