## Supplementary material for "Human iPSC Derived Enamel Organoid Guided by Single-Cell Atlas of Human Tooth Development": File S6_antibodies_and_primers

**File S6: Contains antibodies list and primer lists**

List of the antibodies used for the Immunohistochemistry/Immunofluorescence and Whole mount staining

| **Primary antibodies** | | | | |
| --- | --- | --- | --- | --- |
| **Isotype/ Antigen** | **Company** | **Identifier** | **Dilution factor** | **Host species** |
| AMBN | Santa Cruz | sc-271012; RRID: AB_10613795 | 1:50 | Mouse |
| AMELX | Santa Cruz | sc-365284; RRID: AB_10843799 | 1:50 | Mouse |
| DSPP | Santa Cruz | sc-73632; RRID: AB_2230660 | 1:50 | Mouse |
| ENAM | Thermo Fisher | PA5-25734; RRID: AB_2543234 | 1:100 | Rabbit |
| VIMENTIN | Cell Signalling | 5741S; RRID: AB_10695459 | 1:200 | Rabbit |
| Human Nuclei | Millipore | MAB1281; RRID:AB_94090 | 1:100 | Rabbit |
| CD146 (MCAM) | Abcam | ab75769; RRID: AB_2143375 | 1:200 | Mouse |
| KRT14 | Thermo Fisher | LL002; RRID: AB_306091 | 1:200 | Mouse |
| KRT5 | Sigma-Aldrich | HPA059479; RRID:AB_2684034 | 1:100 | Rabbit |
| ZO-1 | Invitrogen | 33-9100; RRID: AB_87181 | 1:100 | Mouse |
| anti-GFP | Invitrogen | A-1112; RRID:AB_221569 | 1:500 | Rabbit |
| SP6 | Atlas Antibodies | HPA024516; RRID: AB_10960551 | 1:100 | Rabbit |
| CD144 | BD Biosciences | 555661, RRID: AB_396015 | 1:200 | Mouse |
| DAPI | Thermo Fisher | D1306, RRID: AB_2629482 |  |  |

| **Secondary antibodies** | | | | |
| --- | --- | --- | --- | --- |
| **Isotype/ Antigen** | **Company** | **Identifier** | **Dilution factor** | **Host species** |
| Mouse IgG  (Alexa Flour 488) | Thermo Fisher | A11001; RRID: AB_2534069 | 1:500 | Goat |
| Rabbit IgG  (Alexa Flour 488) | Thermo Fisher | A-32731; RRID:AB_2633280 | 1:500 | Goat |
| Mouse IgG  (Alexa Flour 568) | Thermo Fisher | A-11004; RRID:AB_2534072 | 1:500 | Goat |
| Rabbit IgG  (Alexa Flour 568) | Thermo Fisher | A-11036, RRID:AB_10563566 | 1:500 | Goat |
| Mouse IgG  (Alexa Flour 647) | Thermo Fisher | A32728, RRID:AB_2633277 | 1:500 | Goat |
| Rabbit IgG  (Alexa Flour 647) | Thermo Fisher | A32733, RRID:AB_2633282 | 1:500 | Goat |

List of QRT-PCR primers

| **Primer name** | **Sequence** |
| --- | --- |
| PITX1_F | TCCACCAAGAGCTTCACCTT |
| PITX1_R | CGGTGAGGTTGTTGATGTTG |
| PITX2_3F | GTGTGGACCAACCTTACGGAAG |
| PITX2_3R | CGAAGCCATTCTTGCATAGCTCG |
| KRT14_F | CATGAGTGTGGAAGCCGACAT |
| KRT14_R | GCCTCTCAGGGCATTCATCTC |
| bActin_F | TCCCTGGAGAAGAGCTACG |
| bActin_R | GTAGTTTCGTGGATGCCACA |
| NEST_F | GAAACAGCCATAGAGGGCAA |
| NEST_R | TGGTTTTCCAGAGTCTTCAGTGA |
| OCT4_F | GCTGAAGCTGGAGAAGGAGAAGCTG |
| OCT4_R | CAAGGGCCGCAGCTTACACATGTTC |
| P63_F | TTCTTAGCGAGGTTGGGCTG |
| P63_R | GATCGCATGTCGAAATTGCTC |
| TBX1_1F | CGCAGTGGATGAAGCAAATCGTG |
| TBX1_1R | TTTGCGTGGGTCCACATAGACC |
| Brachyury_1F | TATGAGCCTCGAATCCACATAGT |
| Brachyury_1R | CCTCGTTCTGATAAGCAGTCAC |
| AMBN_2F | TTGAGCCTTGAGACAATGAGAC |
| AMBN_2R | AGACCGTGCATCCACAAAGAA |
